# Efficient gene-environment interaction tests for large biobank-scale sequencing studies

**DOI:** 10.1101/2020.04.28.067173

**Authors:** Xinyu Wang, Elise Lim, Ching-Ti Liu, Yun Ju Sung, Dabeeru C. Rao, Alanna C. Morrison, Eric Boerwinkle, Alisa K. Manning, Han Chen

## Abstract

Complex human diseases are affected by genetic and environmental risk factors and their interactions. Gene-environment interaction (GEI) tests for aggregate genetic variant sets have been developed in recent years. However, existing statistical methods become rate limiting for large biobank-scale sequencing studies with correlated samples. We propose efficient Mixed-model Association tests for GEne-Environment interactions (MAGEE), for testing GEI between an aggregate variant set and environmental exposures on quantitative and binary traits in large-scale sequencing studies with related individuals. Joint tests for the aggregate genetic main effects and GEI effects are also developed. A null generalized linear mixed model adjusting for covariates but without any genetic effects is fit only once in a whole genome GEI analysis, thereby vastly reducing the overall computational burden. Score tests for variant sets are performed as a combination of genetic burden and variance component tests by accounting for the genetic main effects using matrix projections. The computational complexity is dramatically reduced in a whole genome GEI analysis, which makes MAGEE scalable to hundreds of thousands of individuals. We applied MAGEE to the exome sequencing data of 41,144 related individuals from the UK Biobank, and the analysis of 18,970 protein coding genes finished within 10.4 CPU hours.

## 1. INTRODUCTION

The variation of traits among individuals in a population results from genetic and environmental factors, as well as their interactions. Gene-environment interaction (GEI) studies can improve our understanding on the biological mechanisms of complex diseases and lead to discoveries of novel genetic associations with effects that vary with environmental exposures (Mcallister et al., 2017). The advancements in next-generation sequencing technologies over the past decade have enabled the increasing availability of large-scale data of low-frequency and rare genetic variants (with minor allele frequency (MAF) 1–5% and <1%, respectively). The single-variant tests conventionally used for testing GEI with common variants are underpowered for rare variants. For example, statistical power for testing GEI with a binary environmental exposure depends on the minor allele counts in both exposed and unexposed groups. Although computationally efficient GEI tests for biobank-scale studies have been developed recently in the context of single-variant tests on unrelated individuals (Bi et al., 2019), critical methodological bottlenecks still exist to expand the sample size and scope of rare variant GEI analyses in large biobank-scale sequencing studies. To increase power for rare variants, various set-based methods have been developed to collapse variants in a particular gene or functional region to investigate how variants in a set affect a phenotype synergistically (Chen et al., 2019; Lee, Wu, & Lin, 2012; Pan, Kim, Zhang, Shen, & Wei, 2014; Sun, J., Zheng, & Hsu, 2013), and to demonstrate whether genetic associations with the phenotype are modified by environment factors in GEI studies (Chen, Meigs, & Dupuis, 2014; Lin et al., 2016; Su, Y., Di, & Hsu, 2017). For example, rareGE is a software tool for GEI tests on rare variants (Chen et al., 2014) that implements three variance component tests: two GEI tests that treat the genetic main effects either as fixed or random effects, and a joint test for the genetic main effects and GEI. (Lin et al., 2016) proposed the interaction sequence kernel association test (iSKAT), which extends the SKAT-O test (Lee et al., 2012) for genetic main effects to a GEI test for rare variants that estimates the genetic main effects using a ridge regression model (Hastie, Tibshirani, & Friedman, 2009). Mixed effects Score Tests for interaction (MiSTi) proposed by (Su et al., 2017) is an extension to MiST (Sun et al., 2013) in the context of GEI.

Most of the aforementioned variant set-based GEI tests were developed to analyze unrelated samples and directly applying these methods to related samples will lead to invalid statistical inference and inflated type I error rates. On the other hand, extending these methods by adding a random effects term to account for relatedness in the generalized linear mixed model (GLMM) framework would result in an intensive computational complexity of *O*(*N*^3^) for each variant set, where *N* is the sample size (Lim, Chen, Dupuis, & Liu, 2020; Mazo Lopera, Coombes, & de Andrade, 2017). For large biobank-scale sequencing studies, such as the UK Biobank, related samples are often present. Therefore, there is an urgent need for computationally efficient variant set-based GEI tests that are scalable to hundreds of thousands of related samples.

Joint tests for genetic main effects and GEI effects are used to identify novel associations previously missed in genetic main effect tests, by accounting for heterogeneous genetic effects in samples with different environmental exposures (Cornelis et al., 2012). Joint tests for common variants have been developed (Chen et al., 2014; Kraft, Yen, Stram, Morrison, & Gauderman, 2007; Selinger-Leneman, Genin, Norris, & Khlat, 2003; Sun, R., Carroll, Christiani, & Lin, 2018), and a variant set-based joint test has been implemented in rareGE (Chen et al., 2014). However, the rareGE joint test relies on a Monte Carlo method to compute the *p* values, which changes slightly with the random number seed used in the analysis, and an analytical solution is not currently available.

We propose a computationally efficient method, Mixed-model Association test for GEne-Environment interactions (MAGEE), to test GEI effects for rare variants that can reduce the computational complexity for testing each variant set from *O*(*N*^3^) to at most *O*(*N*^2^) for related samples, where *N* is the sample size. For samples with well-defined family structures the use of a block diagonal correlation structure can greatly reduce the complexity to *O*(*nN*), where *n* is the maximum number of individuals in each block. For unrelated samples, the computational complexity for testing each variant set in MAGEE is *O*(*N*), with statistical power being close to the existing methods such as rareGE and MiSTi. We also propose analytical joint tests in MAGEE that do not require Monte Carlo approaches for *p* value calculations.

## 2. MATERIALS AND METHODS

### 2.1 Generalized linear mixed models (GLMMs) and main effect tests

We developed two GEI tests and three joint tests based on GLMMs within the MAGEE framework. The full model of MAGEE is:

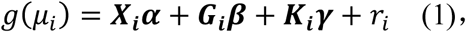

where *g*(·) is the link function of *μ*_*i*_, and *μ*_*i*_ is the conditional mean of the phenotype for individual *i* given covariates ***X***_***i***_, genotypes ***G***_***i***_ and a random intercept *r*_*i*_. ***X***_***i***_ is a row vector of *p* covariates including an intercept, ***G***_***i***_ is a row vector of *q* variants, and ***K***_***i***_ is a row vector of *cq* pairwise GEI terms for *c* environmental factors (which are a subset of the *p* covariates in ***X***_***i***_) and *q* variants. Accordingly, ***α*** is a *p* × 1 vector for the covariate effects, ***β*** is a *q* × 1 vector for the genetic main effects, and ***γ*** is the *cq* × 1 vector for GEI effects. The length *N* vector for the random intercept 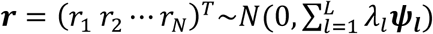, where *λ*_*l*_ are the variance component parameters for *L* random effects, and ***ψ***_***l***_ are *N* × *N* known relatedness matrices.

The genetic main effect model assuming no GEI (***γ*** = **0**) is

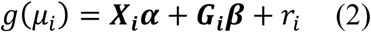

Tests for *H*_0_: ***β*** = **0** can be constructed in the SMMAT framework (Chen et al., 2019). If ***β*** are treated as random effects, the main effect variance component (MV) test is then SKAT (Wu et al., 2011) for related samples (Chen, Meigs, & Dupuis, 2013), from which we can acquire a *p* value *p*_*MV*_. Another main effect test, the main effect hybrid test using Fisher’s method (Fisher, 1928) (MF), which combines the burden test and SKAT, is the efficient hybrid test SMMAT-E (Chen et al., 2019). In this test, we get two *p* values, one from the burden test, *p*_*B*_, and the other from the adjusted SKAT test, *p*_*AS*_. SMMAT-E (Chen et al., 2019) combines these two *p* values to calculate a *p* value for the MF test using Fisher’s method 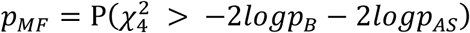.

### 2.2 Interaction tests

In model (2), the score vector for ***γ*** is 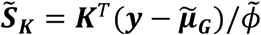, where ***y*** is an *N* × 1 vector of the observed phenotype values, 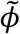 is the dispersion parameter estimate, 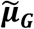 is a vector of fitted values, and 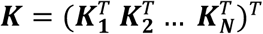 is the *N* × *cq* matrix for the GEI terms. Generally, testing for GEI *H*_0_: ***γ*** = **0** requires adjusting for genetic main effects. Therefore, we need to refit model (2) for every set of genetic variants in a whole genome GEI analysis. To reduce the computational burden, we first fit a global null model without any genetic main effects:

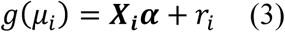

From this model, we can construct score vectors 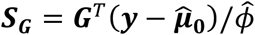 and 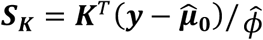 for genetic main effects and GEI effects, respectively, where 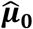 is a vector of fitted values from model (3), and 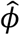 is the dispersion parameter estimate from model (3). Assuming the main effect of genetic variants ***β*** are small, we then approximate the score vector for GEI effects by 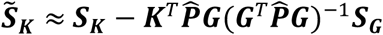 (in appendix A), where 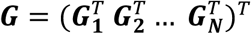 is a *N* × *q* matrix of genetic variants, 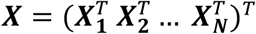 is a *N* × *p* matrix of covariates, 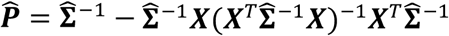 is an *N* × *N* projection matrix, where 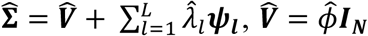 for continuous traits and 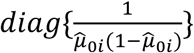 for binary traits, which we estimate from model (3). Using this approximation, in the interaction variance component (IV) test, we assume ***γ*** ∼ *N*(0, *τ****W***_***K***_^2^), where ***W***_***K***_ is a *cq* × *cq* predefined diagonal weight matrix for GEI. Testing for GEI *H*_0_: ***γ*** = **0** is then equivalent to testing the variance component parameter *H*_0_: *τ* = 0, with a test statistic

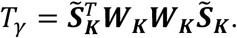

Under the null hypothesis, *T*_*γ*_ asymptotically follows the distribution of 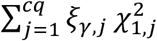, where 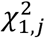 are independent chi-square distributions with 1 degree of freedom (df), and *ξ*_*γ,j*_ are the eigenvalues of ***W***_***K***_**Λ*W***_***K***_, where 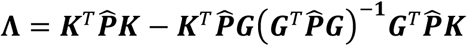 (in appendix A).

Alternatively, we develop the interaction hybrid test using Fisher’s method (IF) to combine a burden-type test and an adjusted variance component test that are asymptotically independent, to achieve superior power than the IV test when the true mean of interaction effects ***γ*** is not close to 0. Specifically, in the IF test we assume 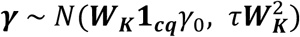, where **1**_***cq***_ is a vector of 1’s with length *cq*, and testing for GEI *H*_0_: ***γ*** = **0** is equivalent to testing *H*_0_: *γ*_0_ = *τ* = 0 simutaneously. We decompose this test into two tests (Chen et al., 2019; Sun et al., 2013). In the first test, we assume *τ* = 0 and test *H*_0_: *γ*_0_ = 0 using the burden score constructed from the global null model (3): 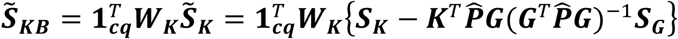. The test statistic

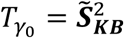

follows a distribution of 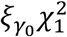 under the null hypothesis *H*_0_: *γ*_0_ = 0, where the scalar 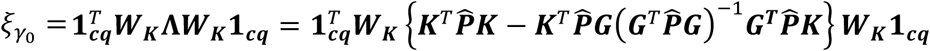.

In the second test, we do not make an assumption that *γ*_0_ = 0 but we assume its true value is small and we test *H*_0_: *τ* = 0 using adjusted scores accounting for the burden effect 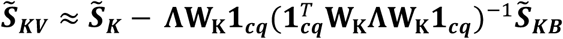 (in appendix B). Then the test statistic can be constructed as:

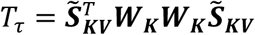

and it asymptotically follows a distribution of 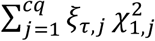 under the null hypothesis *H*_0_: *τ* = 0, where *ξ*_*τ,j*_ are eigenvalues for ***W***_***K***_**Λ**_***KV***_***W***_***K***_, and 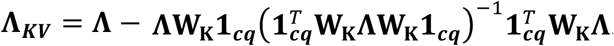.

As 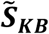 and 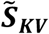 are asymptotically independent (in appendix B), we combine *p* values from the two tests 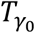 and *T*_*τ*_ using Fisher’s method to compute 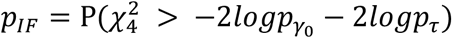, where 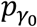 and *p*_*τ*_ are *p* values from the burden-type test *H*_0_: *γ*_0_ = 0 (under the assumption *τ* = 0) and the adjusted variance component test *H*_0_: *τ* = 0, respectively.

### 2.3 Joint tests

As the score vector ***S***_***G***_ for genetic main effects and the adjusted score vector 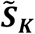 for GEI effects are asymptotically normal with covariance

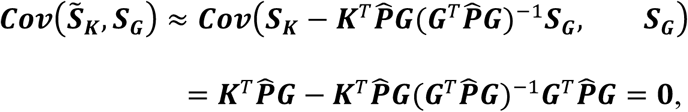

they are asymptotically independent and so are the main effect tests and interaction tests derived using ***S***_***G***_ and 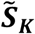, respectively. Therefore, an analytical form of the joint variance component (JV) test for genetic main effects and GEI effects can be constructed by combining *p* values of MV (*p*_*MV*_) and IV (*p*_*IV*_) tests using Fisher’s method as 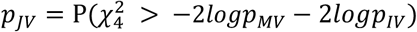, similar to the IF test above.

We can also construct the joint test using MF and IF since they are also derived using ***S***_***G***_ and 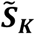 respectively, and both 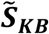 and 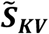 are linear functions of 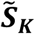 that are asymptotically independent. The four *p* values 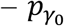 from the GEI burden-type test, *p*_*τ*_ from the GEI adjusted variance component test, as well as the aforementioned components in the genetic main effects MF test, *p*_*B*_ and *p*_*AS*_ – are asymptotically mutually independent. Therefore, the joint hybrid test using Fisher’s method (JF) can be constructed by adding these four *p* values on the log scale, which follows a chi-square distribution with 8 df under the null hypothesis of no genetic main effects or GEI effects (Fisher, 1928) 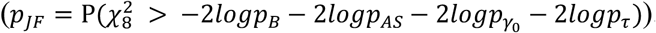. Alternatively, we can apply Fisher’s method to combine two *p* values from MF and IF tests, which follows a chi-square distribution with 4 df under the null hypothesis of no genetic main effects or GEI effects 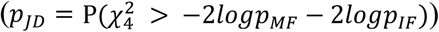. This is a joint hybrid test using double Fisher’s procedures (JD), since each of MF and IF *p* values is already computed using Fisher’s method (refer to Table S1 of the supporting information for a summary of all five new tests: GEI tests IV and IF, and joint tests JV, JF and JD).

### 2.4 Simulations

We conducted extensive simulations to 1) investigate MAGEE’s type I error control in unrelated and related samples; 2) compare *p* values from MAGEE and the existing methods rareGE and MiSTi on the scale of 10^−5^ to 10^−15^ in unrelated samples; and 3) compare the power of each test within the MAGEE framework. All the simulation scenarios were performed for both quantitative and binary traits.

#### 2.4.1 Type I error in unrelated samples

HAPGEN2 (Su, Z., Marchini, & Donnelly, 2011) was used to simulate 200,000 haplotypes on chromosome 22 based on 1000 Genomes project CEU data (Sabeti, 2015) as the reference panel. We randomly paired them into genotypes of 100,000 unrelated individuals. We simulated 40 genotype replicates, and each genotype replicate has a total of 119,317 variants, which were assigned to 1,194 variant sets. The first 1,193 groups had 100 variants per group, while the last group contained 17 variants. The MAF for each variant ranged from 5.0 × 10^−6^ to 0.5, with a mean of 0.18.

We simulated 1,000 phenotype replicates for each genotype replicate. The quantitative trait for individual *i* was simulated from

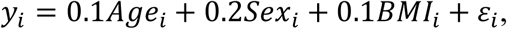

where *Age*_*i*_∼*N*(50, 5), *Sex*_*i*_ ∼ *Bernoulli*(0.5), *BMI*_*i*_ ∼ *N*(25, 4), and the random error *ε*_*i*_ ∼ *N*(0, 1). The binary traits for unrelated sample were simulated as a cohort study from

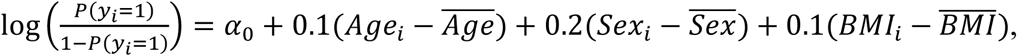

where *y*_*i*_ is the observed phenotype value (either 0 or 1), *Age, Sex*, and body mass index (*BMI*) follow the same distribution as for the quantitative traits, and 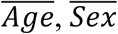, and 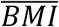 are the mean values for *Age, Sex*, and *BMI*, respectively, and *α* was set to 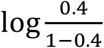. For both quantitative and binary traits, we tested for gene-BMI interactions using beta density weight function with parameters 1 and 25 on the MAF (Wu et al., 2011) for both common and rare genetic variants, so that rare variants would have larger weights than common variants.

#### 2.4.2 Type I error in related samples

Since MAGEE can be applied to both unrelated and related samples, we also simulated 40 genotype replicates for 25,000 families with two parents and two children with a theoretical kinship matrix of 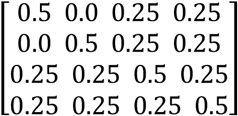, totaling 100,000 individuals. The 100,000 haplotypes for the 50,000 founders were randomly sampled and paired into genotypes, and each child inherited one random haplotype from each parent within the same family. The total number of variants were also 119,317 and were assigned to 1,194 variant sets.

We simulated 1,000 phenotype replicates for each genotype replicate. The quantitative trait for individual *j* in family *i* was simulated from

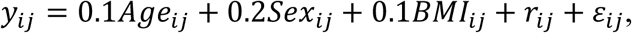

where *Age*_*ij*_ ∼ *N*(50, 5) for parents and *Age*_*ij*_ ∼ *N*(20, 5) for children, *Sex*_*ij*_ ∼ *Bernoulli*(0.5) for both parents and children, BMI for family *i* 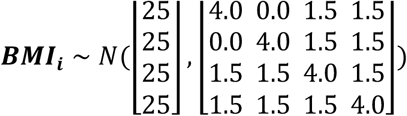, by assuming a heritability of 0.75(Elks et al., 2012), the random effects for family *i* 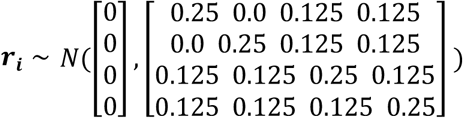, and the random error *ε*_*ij*_ ∼ *N*(0, 0.75).

The binary traits for related sample were simulated as a cohort study from

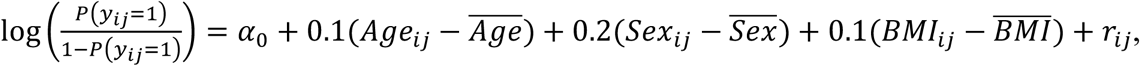

where *Age, Sex, BMI*, and the random effects for family *i* ***r***_***i***_ follow the same distribution as for the quantitative traits, and 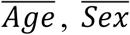, and 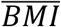 are the population mean values for them, respectively, *α* is 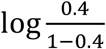. For both quantitative and binary traits, we tested for gene-BMI interactions using beta density weight function with parameters 1 and 25 on the MAF (Wu et al., 2011) for both common and rare genetic variants.

#### 2.4.3 Comparison of *p* values

We simulated both quantitative and binary traits with unrelated individuals to compare the *p* value estimations for MAGEE with existing methods rareGE and MiSTi. Quantitative traits were simulated from 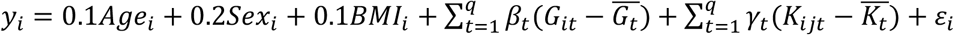, and binary traits were simulated from 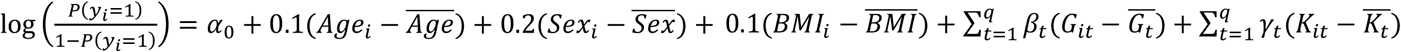, where *G*_*it*_ denoted the *t*-th genetic variant for the *i*-th individual, and 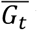 was the mean of the t-th variant, 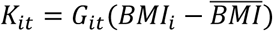 was the GEI term for the *t*-th variant with BMI for individual *i*, and 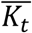 was the mean of the *t*-th interaction term. We conducted tests with sample sizes 2,000, 5,000, and 10,000. In scenario 1, we simulated both genetic main effects *β*_*t*_ and GEI effects *γ*_*t*_ from *clog*_10_(*MAF*_*t*_), with different values of constant *c*. In scenario 2, we simulated only genetic main effects *β*_*t*_ from *clog*_10_(*MAF*_*t*_), and set GEI effects *γ*_*t*_ = 0 so that it was a null hypothesis simulation for interaction tests but an alternative hypothesis simulation for joint tests. The specific values for constant *c* in each scenario and a summary of the simulation scenarios can be found in Table S2. In all these scenarios, *Age*_*i*_, *Sex*_*i*_, *BMI*_*i*_ and *ε*_*i*_ followed the same distribution as in the null simulations for unrelated samples. We simulated one genotype replicate with 100 variants for 10,000 unrelated individuals, and 1,000 phenotype replicates in each scenario. For *β*_*t*_ and *γ*_*t*_, we randomly assigned 80% of them to 0, 10% to have a positive effect, and 10% to have a negative effect.

#### 2.4.4 Power

To investigate the power of each test within the MAGEE framework, we simulated six scenarios using related individuals with sample sizes 20,000, 50,000, and 100,000 in each scenario. In scenarios 1-3, we randomly assigned 80% of genetic main effects *β*_*t*_ and GEI effects *γ*_*t*_ to 0, 10% of *β*_*t*_ and *γ*_*t*_ to have a positive effect, and the remaining 10% of each of *β*_*t*_ and *γ*_*t*_ to have a negative effect. These proportions were changed to 80% null, 16% positive, and 4% negative in scenarios 4-6. In all of the six scenarios, *Age*_*ij*_, *Sex*_*ij*_, *BMI*_*ij*_, *r*_*ij*_, and *ε*_*ij*_ followed the same distributions as the null simulations for related individuals. We simulated one genotype replicate with 100 variants for 100,000 related individuals with the same family structure as the null simulations for related individuals, and 1,000 phenotype replicates in each scenario. Quantitative traits were simulated from 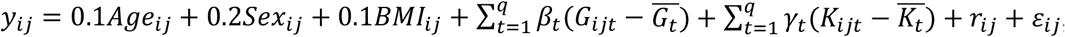, and binary traits were simulated from 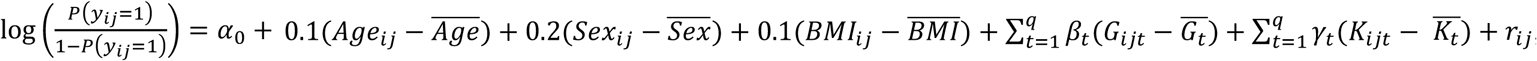, where *G*_*ijt*_ was the *t*-th genetic variant for individual *j* in family *i*, and 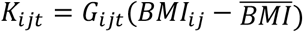 was the GEI term for the *t*-th variant with BMI for individual *j* from family *i*. In scenarios 1 and 4, we simulated both genetic main effects *β*_*t*_ and GEI effects *γ*_*t*_ from *clog*_10_(*MAF*_*t*_), with different values of constant *c*. In scenario 2 and 5, we simulated only genetic main effects *β*_*t*_ from *clog*_10_(*MAF*_*t*_), and set GEI effects *γ*_*t*_ = 0. In scenario 3 and 6, we set genetic main effects *β*_*t*_ = 0, and simulated GEI effects *γ*_*t*_ from *clog*_10_(*MAF*_*t*_). The detailed simulation settings and choices for constant *c* in each scenario for both quantitative and binary trait are listed in Table S3.

### 2.5 Application to UK Biobank whole exome sequencing data

We used the first tranche of UK Biobank whole exome sequencing (WES) data released in March 2019 with 49,959 participants, which included 43,386 White British individuals with non-missing age. We tested for gene-sex interactions on a quantitative trait BMI, as well as a dichotomized trait obesity, which was defined as BMI ≥ 30 (24.1% of the study samples). After removing samples with missing genetic ancestry or gender and sex mismatches, the total number of individuals was 41,144, including 18,925 men and 22,219 women. For BMI, we fit a heteroscedastic linear mixed model (Conomos et al., 2016) that allowed the residual variance to be different in men and women, adjusting for sex, age, age^2^, the interaction terms between sex and age, age^2^, and the top ten ancestry principal components (PCs). For obesity, we adjusted for the same covariates in a GLMM. We used the population-level genotype data generated from the Functionally Equivalent (FE) pipeline (Regier et al., 2018). The variant sets were defined as protein-coding regions for the WES data and a total of 18,970 protein-coding regions with 9,760,758 variants were available for analysis. Variants with MAF less than 0.001 were excluded from the analysis. We applied a beta density weight function with parameters 1 and 25 on the MAF (Wu et al., 2011), and performed both GEI and joint tests using a single thread on a computing server.

## 3. RESULTS

### 3.1 Simulation Studies

#### 3.1.1 Type I error

Table 1 summarizes the empirical type I error rates of MAGEE tests for quantitative and binary traits with 100,000 unrelated sample at significance levels of 0.05, 1.0 × 10^−4^, and 2.5 × 10^−6^. In each cell, empirical type I error rates were estimated from 47,760,000 *p* values under the null hypothesis (40 genotype replicates × 1,000 phenotype replicates × 1,194 variant sets) of the null model for unrelated samples. In Figure S1, the quantile-quantile (QQ) plots for both quantitative and binary traits show that all MAGEE main effect tests, GEI tests, and joint tests are well calibrated.

**Table 1.** Empirical type I error rates of MAGEE tests for 100,000 unrelated individuals at significance levels of 0.05, 1.0 × 10^−4^, and 2.5 × 10^−6^.

| Test | Quantitative trait<br>Significance Level |  |  | Binary trait<br>Significance Level |  |  |
| --- | --- | --- | --- | --- | --- | --- |
| | 0.05 | $1.0 \times 10^{-4}$ | $2.5 \times 10^{-6}$ | 0.05 | $1.0 \times 10^{-4}$ | $2.5 \times 10^{-6}$ |
| MV | 0.050 | $1.00 \times 10^{-4}$ | $2.74 \times 10^{-6}$ | 0.050 | $1.02 \times 10^{-4}$ | $2.35 \times 10^{-6}$ |
| MF | 0.050 | $1.01 \times 10^{-4}$ | $2.49 \times 10^{-6}$ | 0.050 | $1.00 \times 10^{-4}$ | $2.84 \times 10^{-6}$ |
| IV | 0.050 | $1.04 \times 10^{-4}$ | $2.47 \times 10^{-6}$ | 0.050 | $1.01 \times 10^{-4}$ | $2.31 \times 10^{-6}$ |
| IF | 0.050 | $1.01 \times 10^{-4}$ | $2.70 \times 10^{-6}$ | 0.050 | $9.93 \times 10^{-5}$ | $2.14 \times 10^{-6}$ |
| JV | 0.050 | $1.02 \times 10^{-4}$ | $2.51 \times 10^{-6}$ | 0.050 | $1.01 \times 10^{-4}$ | $2.44 \times 10^{-6}$ |
| JF | 0.050 | $1.00 \times 10^{-4}$ | $2.55 \times 10^{-6}$ | 0.050 | $1.02 \times 10^{-4}$ | $2.59 \times 10^{-6}$ |
| JD | 0.050 | $1.00 \times 10^{-4}$ | $2.53 \times 10^{-6}$ | 0.050 | $1.01 \times 10^{-4}$ | $2.40 \times 10^{-6}$ |

Similarly, Table 2 shows the empirical type I error rates of MAGEE tests for 100,000 related samples from 25,000 families at the same significance levels as Table 1. Figure 1 shows that with related individuals in the sample, existing methods that ignore the correlation structure, such as rareGE and MiSTi, give inflated type I error rates. In contrast, MAGEE tests control type I errors successfully. Due to the computational speed issue, rareGE and MiSTi do not scale up to the sample size of 100,000. Therefore, analyses in Figure 1 were performed using 10,000 related samples from 2,500 families, in which 119,400 *p* values (1 genotype replicate × 100 phenotype replicates × 1,194 variant sets) were estimated. Figure S2 shows the QQ plots of MAGEE tests for quantitative and binary traits with 100,000 related samples from 47,760,000 *p* values, which are consistent with our results from 100,000 unrelated samples in Figure S1.

**Table 2.** Empirical type I error rates of MAGEE tests for 100,000 related individuals at significance levels of 0.05, 1.0 × 10^−4^, and 2.5 × 10^−6^.

| Test | Quantitative trait<br>Significance Level |  |  | Binary trait<br>Significance Level |  |  |
| --- | --- | --- | --- | --- | --- | --- |
| | 0.05 | $1.0 \times 10^{-4}$ | $2.5 \times 10^{-6}$ | 0.05 | $1.0 \times 10^{-4}$ | $2.5 \times 10^{-6}$ |
| MV | 0.050 | $9.97 \times 10^{-5}$ | $2.32 \times 10^{-6}$ | 0.050 | $9.92 \times 10^{-5}$ | $2.28 \times 10^{-6}$ |
| MF | 0.050 | $1.01 \times 10^{-4}$ | $2.72 \times 10^{-6}$ | 0.050 | $9.93 \times 10^{-5}$ | $1.86 \times 10^{-6}$ |
| IV | 0.050 | $9.82 \times 10^{-5}$ | $1.99 \times 10^{-6}$ | 0.048 | $8.90 \times 10^{-5}$ | $2.07 \times 10^{-6}$ |
| IF | 0.050 | $9.87 \times 10^{-5}$ | $1.97 \times 10^{-6}$ | 0.048 | $8.81 \times 10^{-5}$ | $2.14 \times 10^{-6}$ |
| JV | 0.050 | $9.99 \times 10^{-5}$ | $2.81 \times 10^{-6}$ | 0.049 | $9.55 \times 10^{-5}$ | $2.14 \times 10^{-6}$ |
| JF | 0.050 | $9.98 \times 10^{-5}$ | $2.74 \times 10^{-6}$ | 0.048 | $9.30 \times 10^{-5}$ | $2.11 \times 10^{-6}$ |
| JD | 0.050 | $1.01 \times 10^{-4}$ | $2.83 \times 10^{-6}$ | 0.048 | $9.14 \times 10^{-5}$ | $2.16 \times 10^{-6}$ |

**Figure 1.**
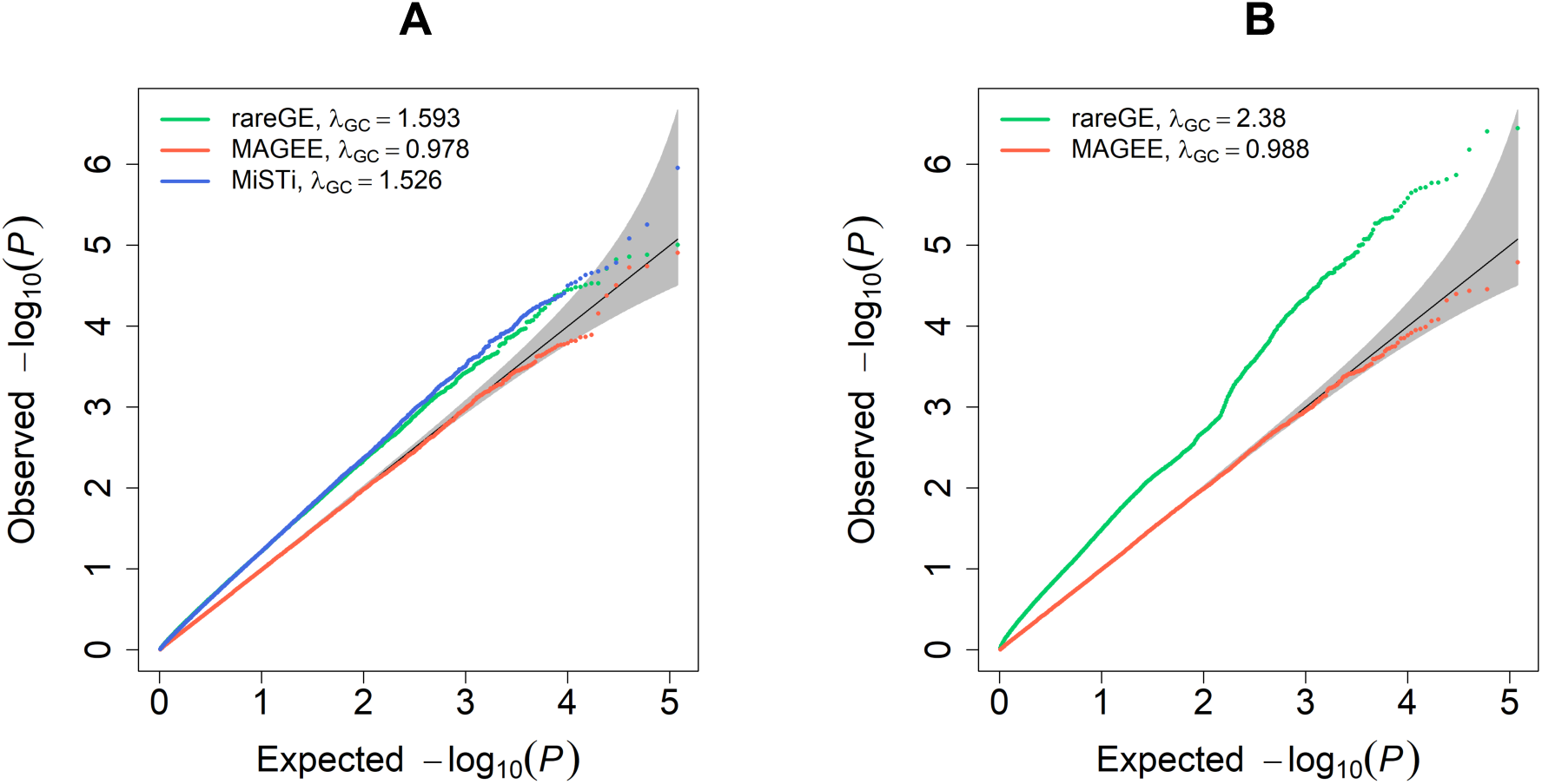
Quantile-Quantile plots of MAGEE, rareGE and MiSTi tests on quantitative traits in 10,000 related samples. (A) MAGEE IV, rareGE and MiSTi GEI tests. (B) MAGEE JV and rareGE joint tests.

#### 3.1.2 Computation time and *p* value comparison

We compared the *p* values from MAGEE with those from rareGE and MiSTi for both quantitative and binary traits in 2,000, 5,000, and 10,000 unrelated samples. Each panel in Figure 2 (scenario 1 for quantitative trait: both genetic main effects and GEI effects) displays 1,000 *p* values from quantitative trait analyses with unrelated samples when both genetic main effects and GEI effects were simulated. MAGEE IV, IF, and JV tests have close *p* values with rareGE GEI test, MiSTi test, and rareGE JOINT test, respectively, and the accuracy increases with the sample size. Similar results are found in Figure S3 (scenario 2 for quantitative trait: genetic main effects only), in which only the genetic main effects but no GEI effects were simulated, as well as Figures S4 and S5 (scenario 1 and scenario 2 for binary traits).

**Figure 2.**
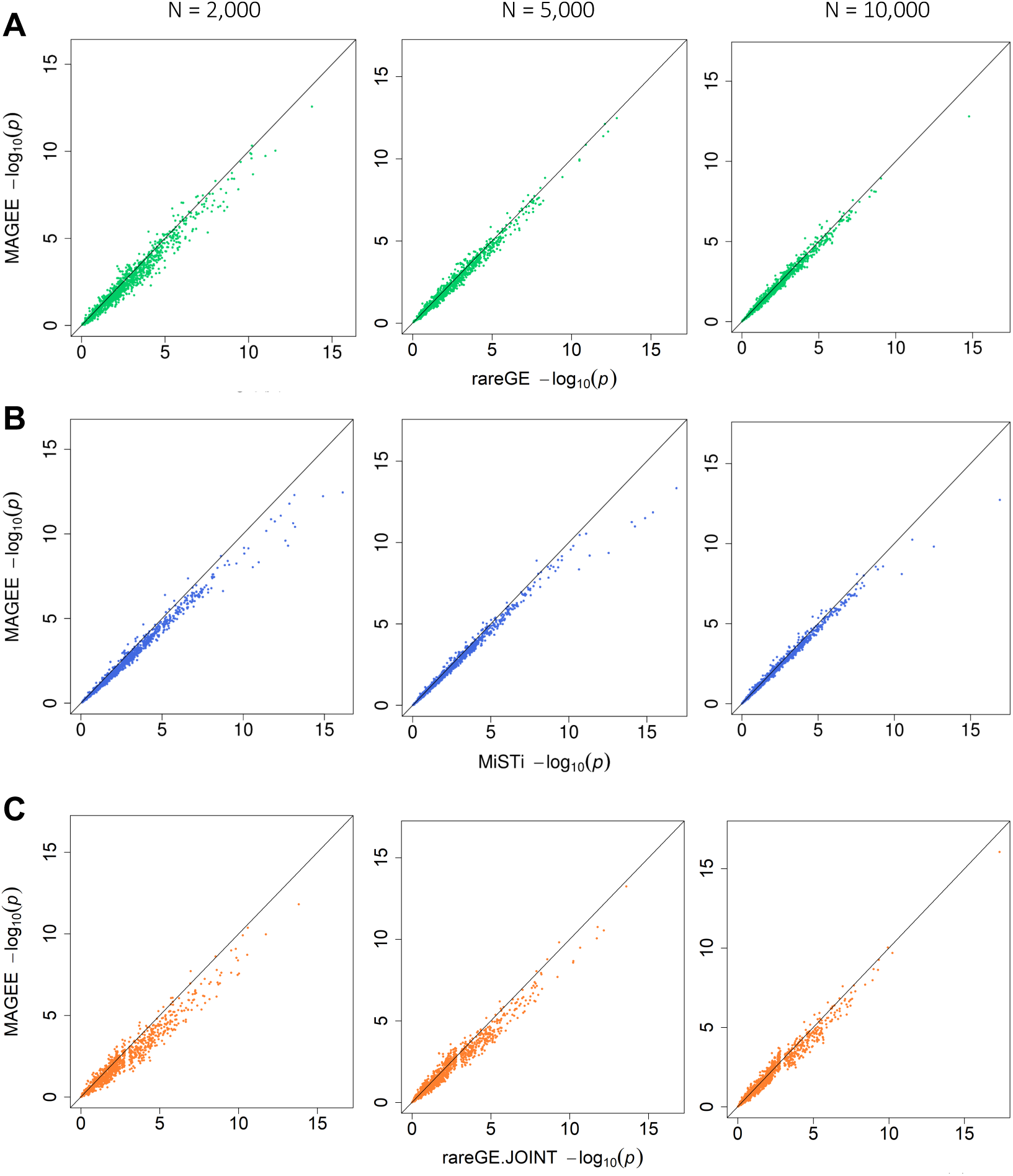
Comparison of *p* values from MAGEE versus rareGE and MiSTi tests on quantitative traits when both genetic and GEI effects were present (scenario 1) in 2,000, 5,000, and 10,000 unrelated samples. (A) MAGEE IV versus rareGE GEI tests. (B) MAGEE IF versus MiSTi GEI tests. (C) MAGEE JV versus rareGE joint tests.

Figure 3 compares the CPU time per *p* value on a single thread for MAGEE, rareGE, and MiSTi in quantitative trait analyses using unrelated individuals. As both rareGE and MiSTi require fitting a separate statistical model for each genetic variant set, their CPU time increases dramatically with the sample size, while MAGEE tests remain computationally efficient in large samples. Similar results are also observed in Figure S6 from binary trait analyses. In addition, MAGEE joint tests are performed simultaneously with the main effect and GEI tests. For example, when performing the JV test, MV and IV test results will be produced automatically; when performing either the JF or JD test, MF and IF test results will also be computed automatically.

**Figure 3.**
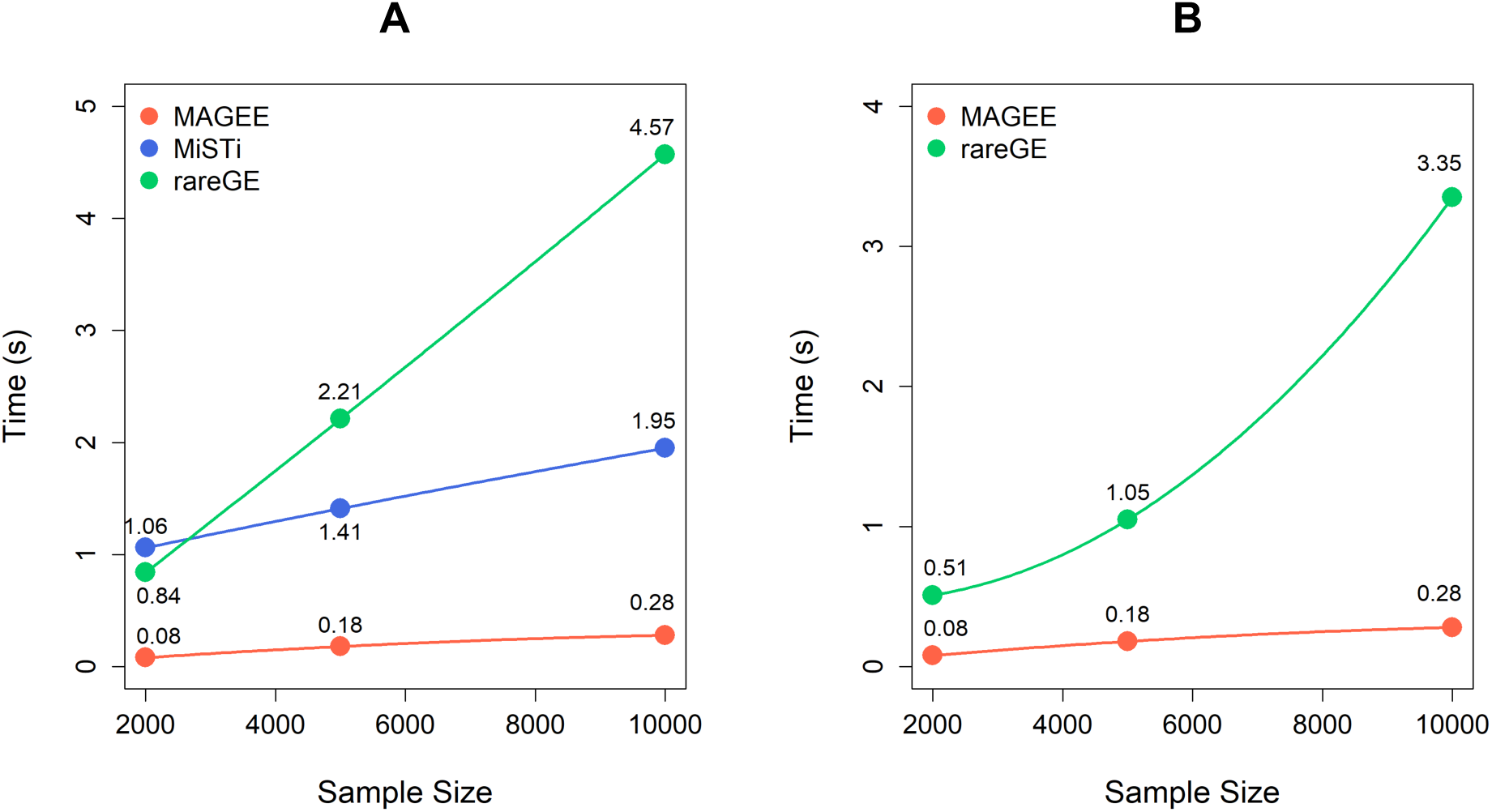
CPU time per *p* value of MAGEE, rareGE and MiSTi tests on quantitative traits in unrelated samples. (A) MAGEE, rareGE and MiSTi GEI tests. (B) MAGEE and rareGE joint tests.

#### 3.1.3 Power

Figure 4 shows the empirical power of the seven tests within the MAGEE framework in analyzing quantitative traits (scenario 1 to 6), at the significance level of 2.5 × 10^−6^ with 20,000, 50,000, and 100,000 related samples, respectively. The top panels show results from the scenarios 1-3, which have variants with 80% null, 10% positive, and 10% negative effects. When both genetic main effects and GEI effects are present (Figure 4A), the three joint tests are most powerful and they have very close power that increases along with the sample size. In the other two scenarios with only either genetic main effects (Figure 4B) or GEI effects (Figure 4C), the joint tests are less powerful than main effects tests or GEI tests, respectively. In general, the variance component tests (MV, IV and JV) and the hybrid tests (MF, IF, JF and JD) have close power, with the hybrid tests power being slightly more powerful in these simulation settings. The bottom panels show results from the scenarios 4-6, which have variants with 80% null, 16% positive, and 4% negative effects. The results are similar with scenarios 1-3 except that the hybrid tests have recognizable higher power compared to the variance component tests, for all main effect, GEI and joint tests. Furthermore, the JF test is slightly less powerful than the JD test, except in Figure 4A and 4D, when both genetic main effects and GEI effects were simulated. Similar results with six simulation scenarios for binary traits are shown in Figure S7. In general settings, we would recommend IF for the GEI test, and JD for the joint test, if we do not have any prior knowledge about the genetic architecture of main effects or GEI effects.

**Figure 4.**
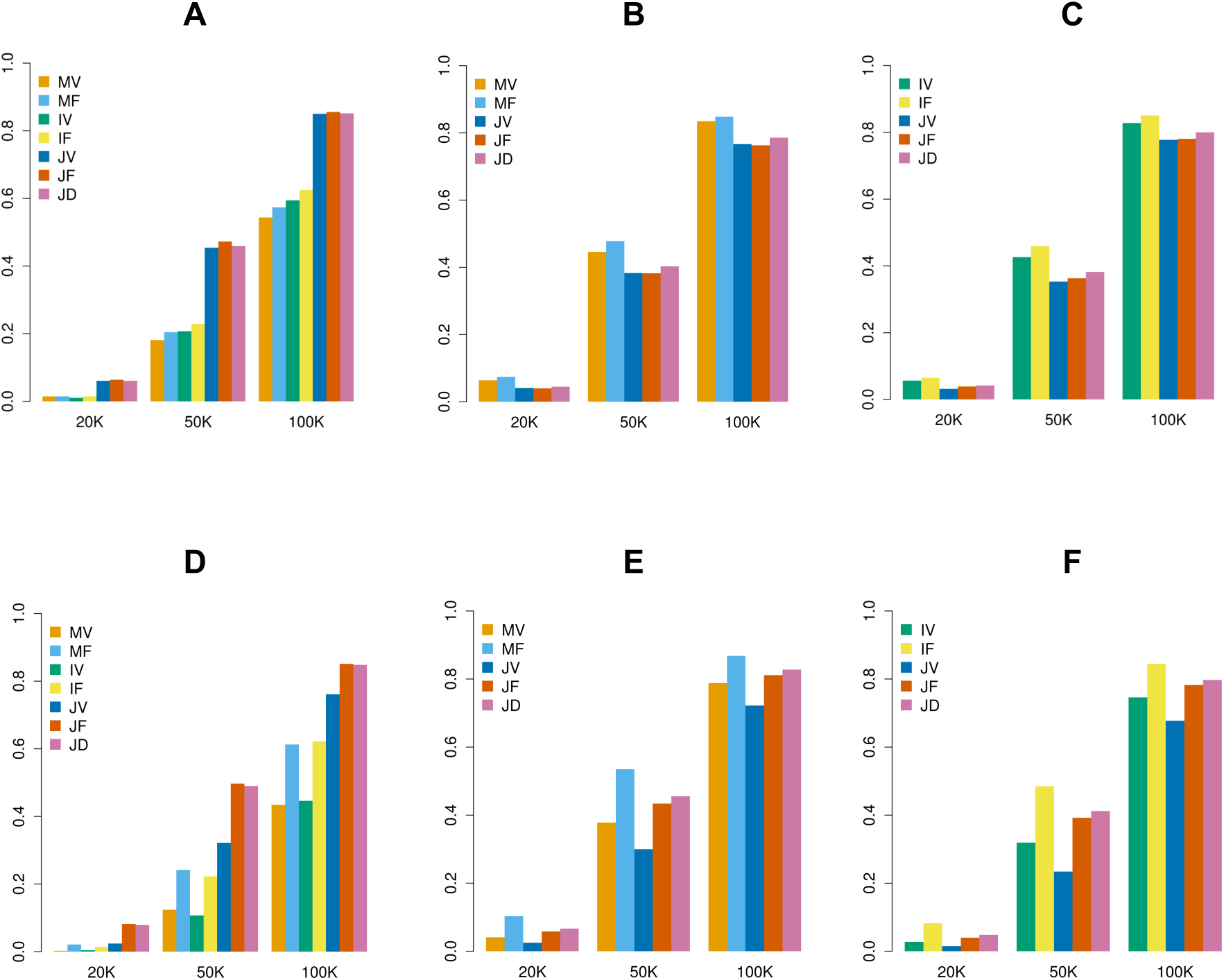
Empirical power of MAGEE tests on quantitative traits in 20,000, 50,000, and 100,000 related samples. (A) Scenario 1: 80% null variants, 10% causal variants with positive effects and 10% causal variants with negative effects for both genetic main effects and GEI effects. (B) Scenario 2: 80% null variants, 10% causal variants with positive effects and 10% causal variants with negative effects for genetic main effects only. (C) Scenario 3: 80% null variants, 10% causal variants with positive effects and 10% causal variants with negative effects for GEI effects only. (D) Scenario 4: 80% null variants, 16% causal variants with positive effects and 4% causal variants with negative effects for both genetic main effects and GEI effects. (E) Scenario 5: 80% null variants, 16% causal variants with positive effects and 4% causal variants with negative effects for genetic main effects only. (F) Scenario 6: 80% null variants, 16% causal variants with positive effects and 4% causal variants with negative effects for GEI effects only.

### 3.2 UK Biobank whole exome sequencing data

The analysis of all 18,970 protein coding regions were finished in 10.4 hours using a single thread on a computing server for a single phenotype. Figure 5 shows that the IF tests are well calibrated for both the quantitative trait BMI and the dichotomized trait obesity, while the mild inflations in the JD tests are likely attributable to the main effect tests, as expected for polygenic traits. However, from Figure 6 one can see that across the 18,970 protein-coding regions, only one significant *p* value in the melanocortin 4 receptor (*MC4R*) gene region on chromosome 18 was found in the JD test of gene-sex interaction for BMI at the significance level of 0.05/18,970 = 2.64 × 10^oú^ after Bonferroni correction for multiple testing (Bland & Altman, 1995). The IF test *p* value in this region was not significant (*p* = 0.26), so the significant signal in the JD test (*p* = 2.29 × 10^−7^) was mostly driven by genetic main effects (MF test *p* = 4.56 × 10^−8^). *MC4R* is one of the most common monogenic cause of severe obesity in humans (Krashes, Lowell, & Garfield, 2016; Vaisse, Clement, Guy-Grand, & Froguel, 1998). Previously, genome-wide association studies have identified common variants in or near *MC4R* with sex-specific effects on human brain structure and eating behavior (Horstmann et al., 2013), but it is unknown whether there are sex-specific genetic effects on BMI in this region.

**Figure 5.**
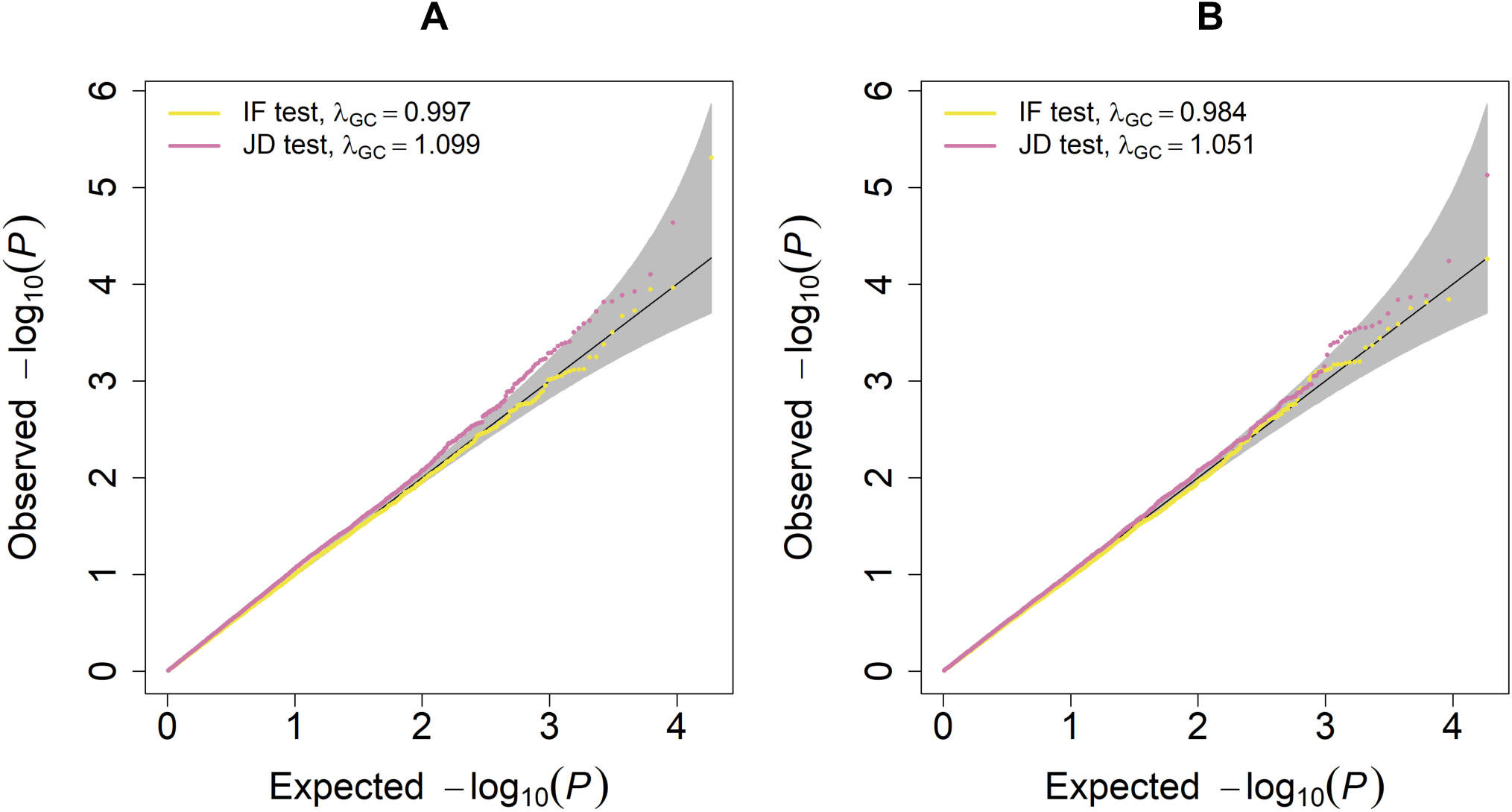
Quantile-Quantile plots of UK Biobank WES data analysis using MAGEE tests for gene-sex interaction effects on BMI and obesity. (A) BMI analysis. (B) Obesity analysis.

**Figure 6.**
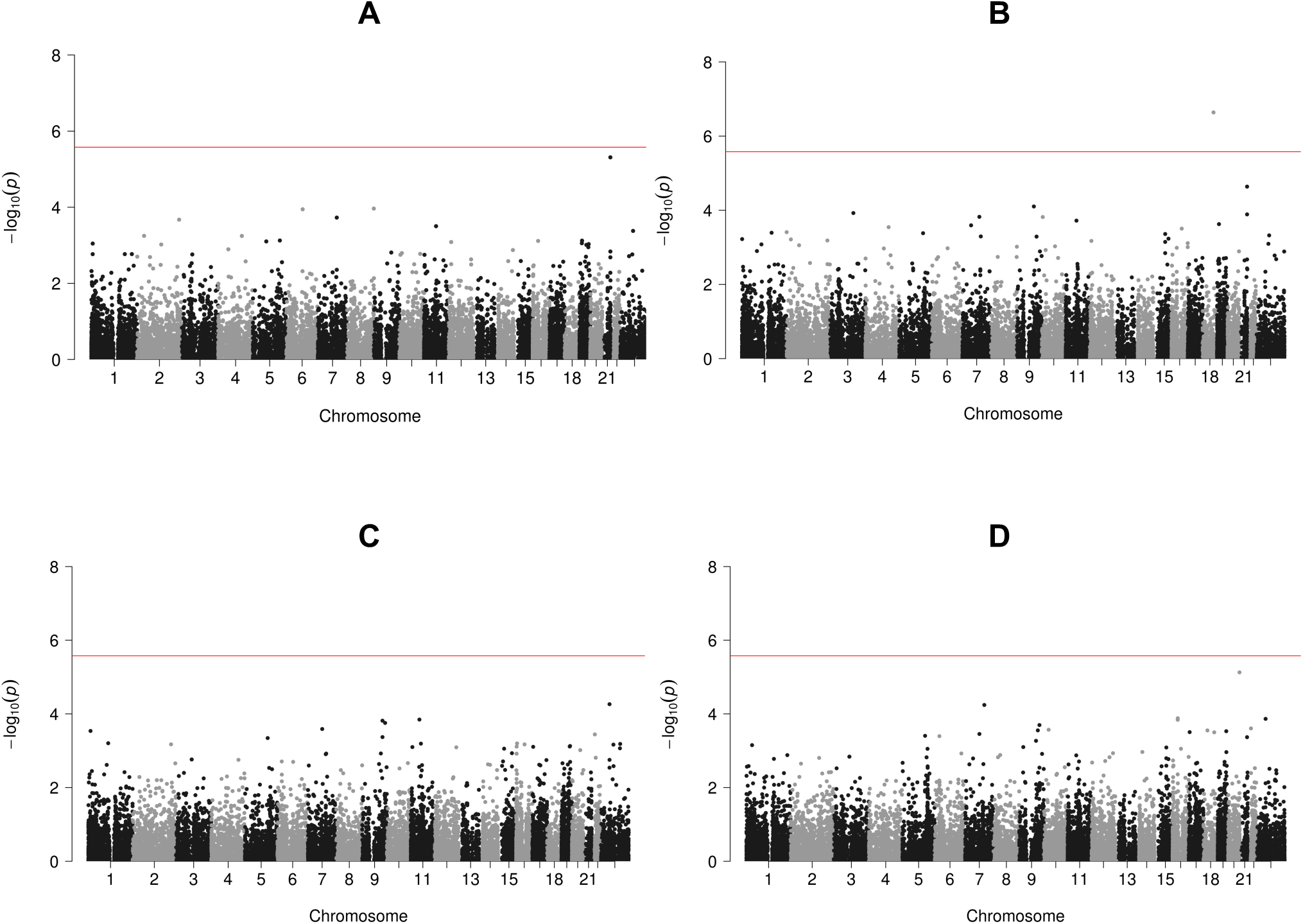
Manhattan plots of UK Biobank WES data analysis using MAGEE tests for gene-sex interaction effects on BMI and obesity. (A) MAGEE IF test on BMI. (B) MAGEE JD test on BMI. (C) MAGEE IF test on obesity. (D) MAGEE JD test on obesity.

## 4. DISCUSSION

We have developed computationally efficient variant set-based mixed model GEI tests and joint tests in the MAGEE framework, which can be applied to both quantitative and binary traits in large biobank-scale sequencing studies with hundreds of thousands of possibly related individuals. We have shown in simulation studies that existing GEI tests for variant sets developed for unrelated samples would have inflated type I error rates when applied to related samples, while MAGEE successfully controls type I errors in both unrelated and related samples. MAGEE requires fitting a global null model only once for all the tests across the whole genome, and it uses two matrix projection approaches to approximate the test statistics. MAGEE accounts for sample relatedness using GLMMs, and it greatly reduces the computational complexity for testing each variant set from *O*(*N*^3^) to no more than *O*(*N*^2^) with highly accurate approximations, making it the method of choice. For samples with a block-diagonal relatedness structure (such as family studies), the computational complexity for testing each variant set is further reduced to *O*(*nN*), where *n* is the maximum number of individuals in each block (e.g., the family size), which is often much smaller than the total sample size *N*. For unrelated samples, MAGEE tests have a computational complexity of *O*(*N*). The CPU time of MAGEE is much smaller than existing methods, since it does not require fitting a separate statistical model to account for genetic main effects, for each variant set in the whole genome analysis. Moreover, the MAGEE joint tests are unique because *p* values are computed analytically, instead of using the Monte Carlo approaches as implemented in rareGE (Chen et al., 2014). Based on our simulation studies, the hybrid test IF is more powerful than the variance component test IV for GEI effects, especially when the interaction effects do not have a mean of 0, and the hybrid test JD is usually the most powerful joint test. When both genetic main effects and GEI effect are present on approximately the same scale, the hybrid test JF is slightly more powerful than the JD test, but the power difference is negligible. In reality, with little knowledge on the genetic architecture of a quantitative or binary trait, the IF test is recommended for identifying GEI effects, and the JD test is recommended for identifying genetic associations that allow for heterogeneous effects in different environmental exposures.

We applied MAGEE to the WES data from the UK Biobank. The results showed that MAGEE *p* values were well calibrated in these real data applications, and we identified an association between BMI and *MC4R* gene from the joint test. However, we did not find any significant *p* values from the interaction test. It is possible that interaction effects may be too small to identify in 41,144 samples. As the WES project is ongoing in the UK Biobank, we hope to revisit gene-environment interaction analyses when WES data from more UK Biobank samples are released in the coming years.

Recently, StructLMM was developed to test GEI effects for a single genetic variant with high-dimensional environmental factors (Moore et al., 2019). While MAGEE can test GEI effects for multiple genetic variants with multiple environmental factors, its performance in a high-dimensional setting (e.g., a large number of genetic variants along with hundreds of environmental factors) has not been fully investigated. Moreover, MAGEE GEI and joint tests using summary statistics in a meta-analysis setting can further boost statistical power by combining association evidence from multi-million samples from large-scale sequencing studies in upcoming years. Our MAGEE framework provides a foundation for future research in these directions.

## ACKNOWLEDGMENTS

This research has been conducted using the UK Biobank Resource under Application Number 42646. The authors acknowledge the Texas Advanced Computing Center (TACC, https://www.tacc.utexas.edu) at The University of Texas at Austin for providing high performance computing (HPC) resources that have contributed to the research results reported within this paper. This work was supported by National Institutes of Health (NIH) grants R00 HL130593 and R01 HL145025.

## SOFTWARE

We have implemented MAGEE in an R package, available at https://github.com/xwang21/magee.

## SUPPORTING INFORMATION

Supporting information includes seven supplemental figures and three supplemental tables.

## APPENDIX A: APPROXIMATIONS FOR THE SCORE VECTOR 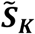

Let 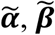, and 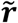 be the estimates for ***α, β***, and ***r*** from Equation 2, 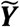 is the working vector for the GLMM and 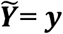 for quantitative trait, and 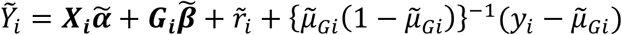 for binary traits.

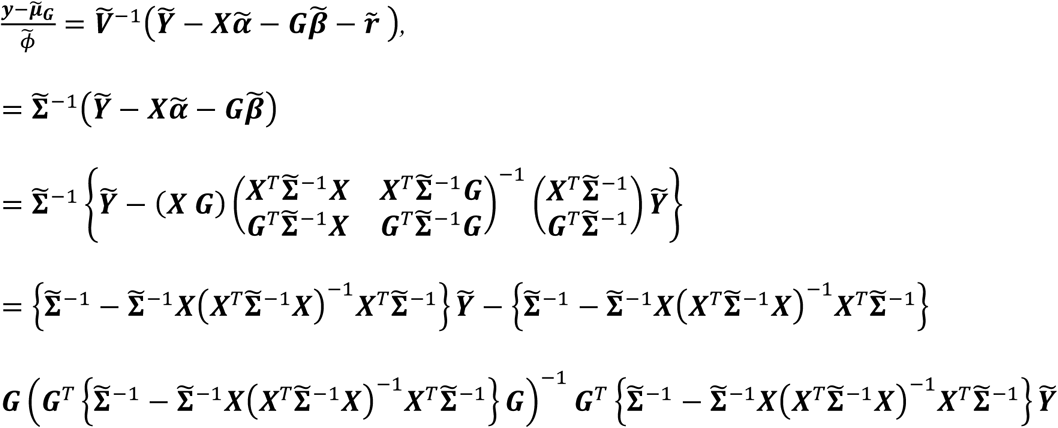

Assuming the true value of ***β*** is small, including the ***G***_***i***_***β*** term in Equation (2) does not dramatically change the variance component estimates for *τ* and *ϕ* from Equation (3), we can approximate that 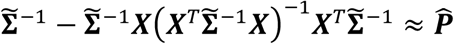 and 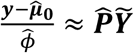, so

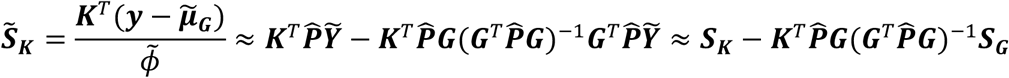

We can rewrite 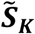 in the matrix form as 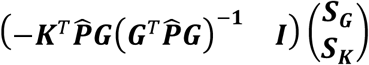, where 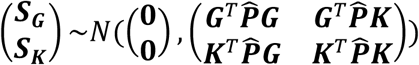 the variance of 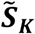 is then

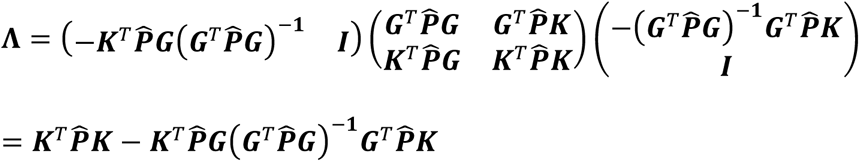

## APPENDIX B: APPROXIMATIONS FOR THE SCORE VECTOR 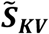

To get 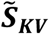, we need to fit Equation (A1)

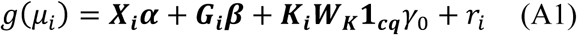

Let 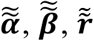, and 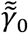 be the estimates for ***α, β, r*** and *γ*_0_ from Equation (A1), 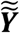 is the working vector for Equation (A1) and 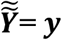 for quantitative trait, and 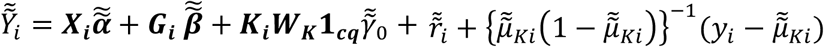 for binary traits, where 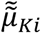 is the mean for individual i after fitting Equation (A1). The mean vector after fitting Equation (A1) is 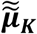.

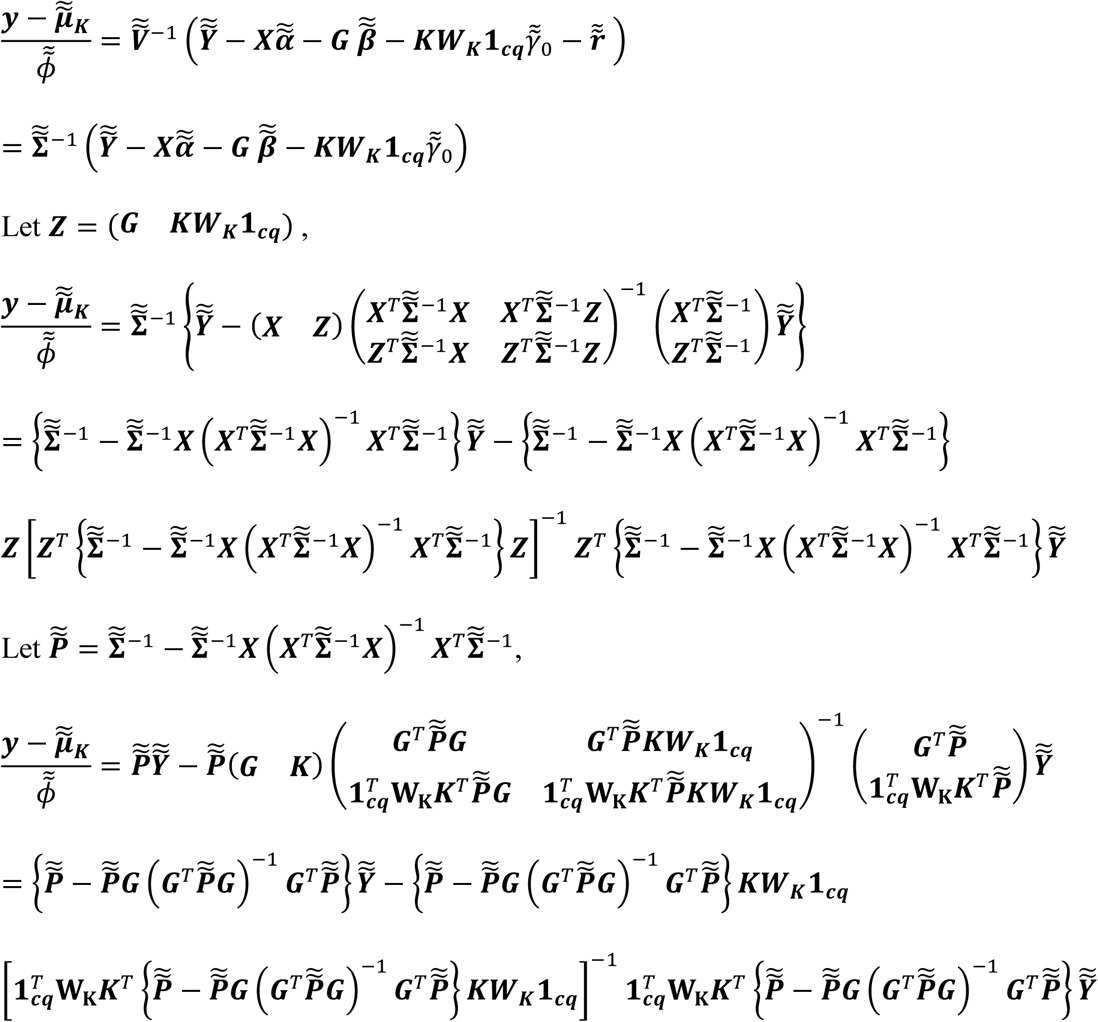

Assuming the true values of ***β*** and *γ*_0_ are small, including the terms ***G***_***i***_***β*** and ***K***_***i***_***W***_***K***_**1**_***cq***_*γ*_0_ in Equation (A1) does not dramatically change the variance component estimates for *τ* and *ϕ* from Equation (3), so we can approximate that 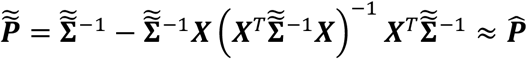 and 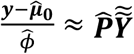, so

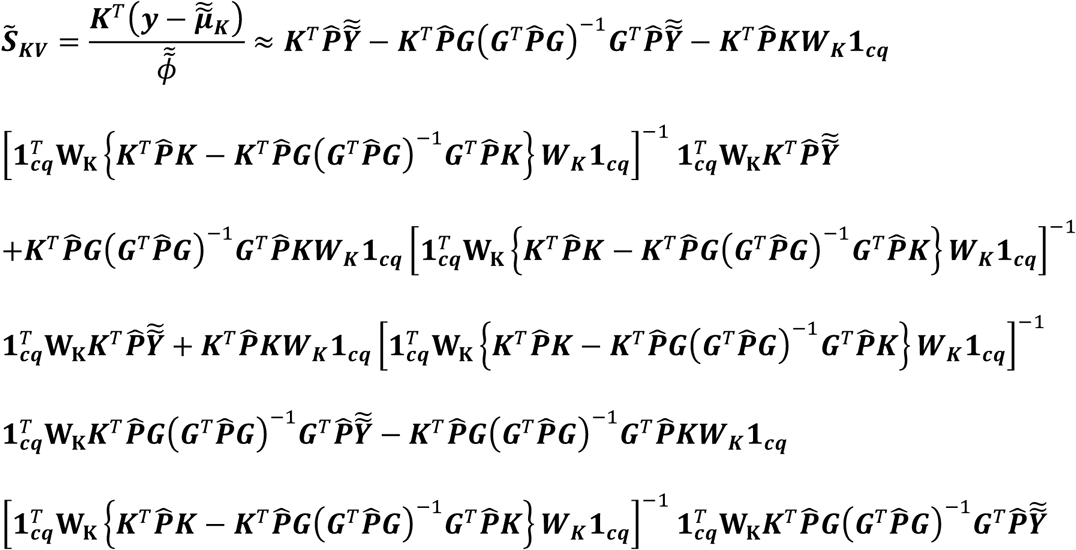

Notice that 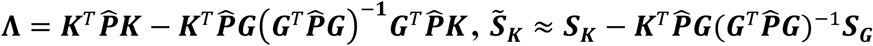 and 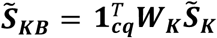,

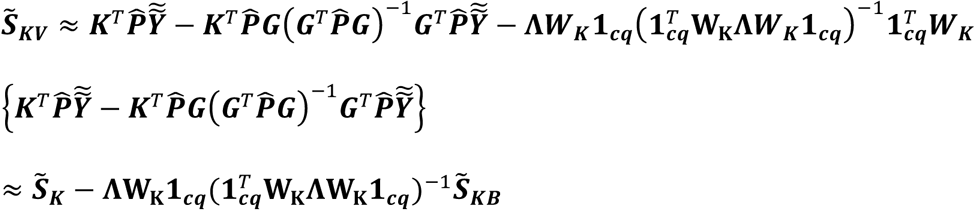

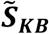 and 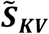 are asymptotically independent because they are asymptotically normal with covariance

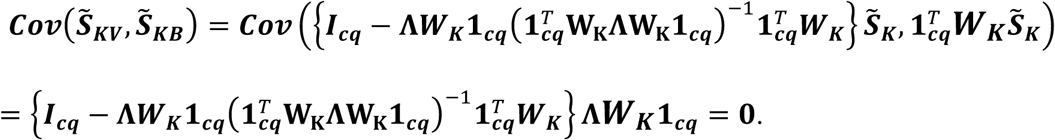

