## Supplemental Figures and Tables for "Efficient gene-environment interaction tests for large biobank-scale sequencing studies"

### SUPPORTING INFORMATION

#### FIGURES

Figure S1. Quantile-Quantile plots of MAGEE tests on quantitative and binary traits in 100,000 unrelated samples under the null model. (A) Main effect (MV and MF) tests on quantitative traits. (B) GEI (IV and IF) tests on quantitative traits. (C) Joint (JV, JF and JD) tests on quantitative traits. (D) Main effect (MV and MF) tests on binary traits. (E) GEI (IV and IF) tests on binary traits. (F) Joint (JV, JF and JD) tests on binary traits.

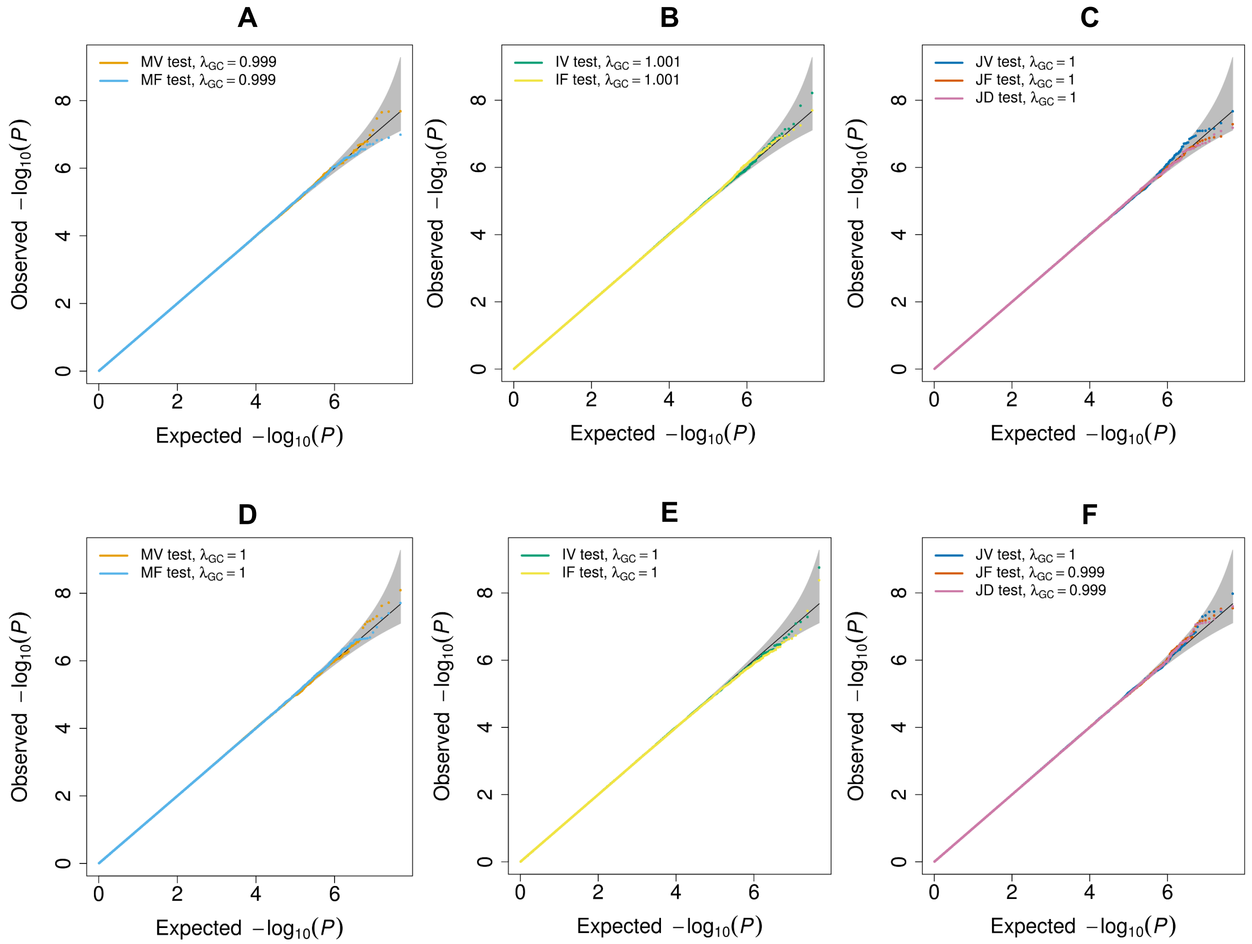

Figure S2. Quantile-Quantile plots of MAGEE tests on quantitative and binary traits in 100,000 related samples under the null model. (A) Main effect (MV and MF) tests on quantitative traits. (B) GEI (IV and IF) tests on quantitative traits. (C) Joint (JV, JF and JD) tests on quantitative traits. (D) Main effect (MV and MF) tests on binary traits. (E) GEI (IV and IF) tests on binary traits. (F) Joint (JV, JF and JD) tests on binary traits.

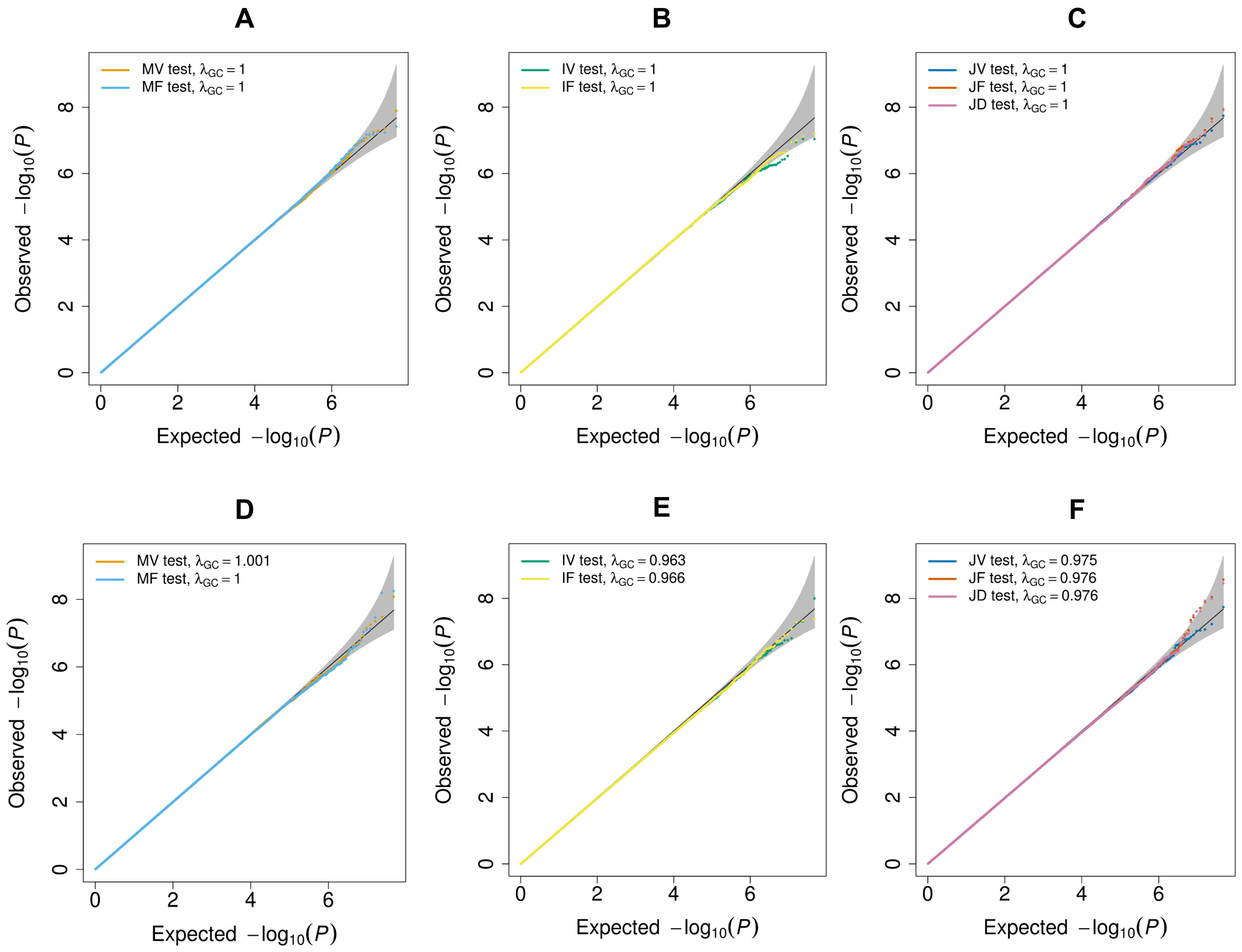

Figure S3. Comparison of *p* values from MAGEE versus rareGE and MiSTi tests on quantitative traits when only genetic effects but not GEI effects were present (scenario 2) in 2,000, 5,000, and 10,000 unrelated samples. (A) MAGEE IV vs. rareGE GEI tests. (B) MAGEE IF vs. MiSTi tests. (C) MAGEE JV vs. rareGE JOINT tests.

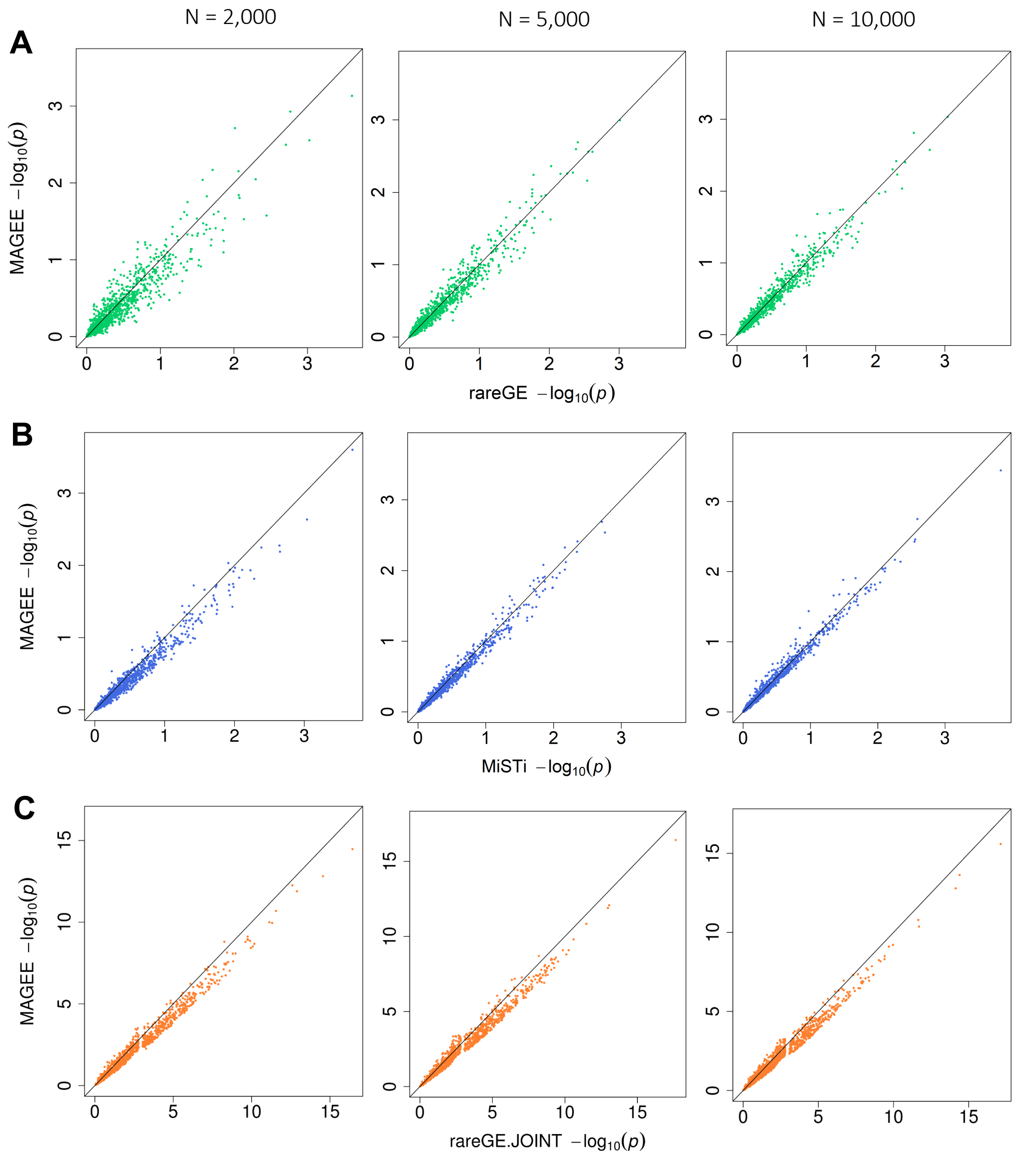

Figure S4. Comparison of *p* values from MAGEE versus rareGE and MiSTi tests on binary traits when both genetic and GEI effects were present (scenario 1) in 2,000, 5,000, and 10,000 unrelated samples. A) MAGEE IV vs. rareGE GEI tests. (B) MAGEE IF vs. MiSTi tests. (C) MAGEE JV vs. rareGE JOINT tests.

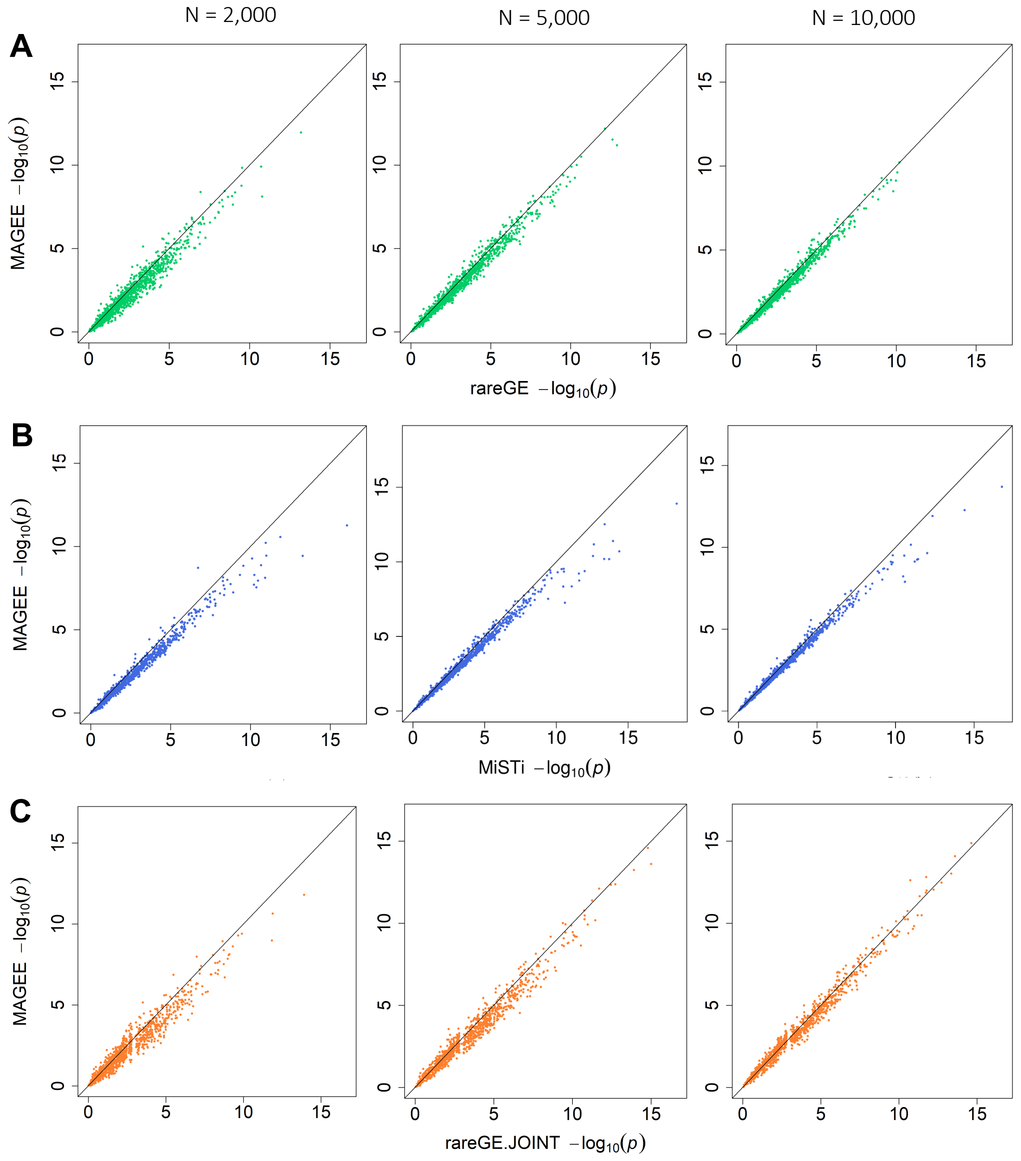

Figure S5. Comparison of *p* values from MAGEE versus rareGE and MiSTi tests on binary traits when only genetic effects but no GEI effects were present (scenario 2) in 2,000, 5,000, and 10,000 unrelated samples. (A) MAGEE IV vs. rareGE GEI tests. (B) MAGEE IF vs. MiSTi tests. (C) MAGEE JV vs. rareGE JOINT tests.

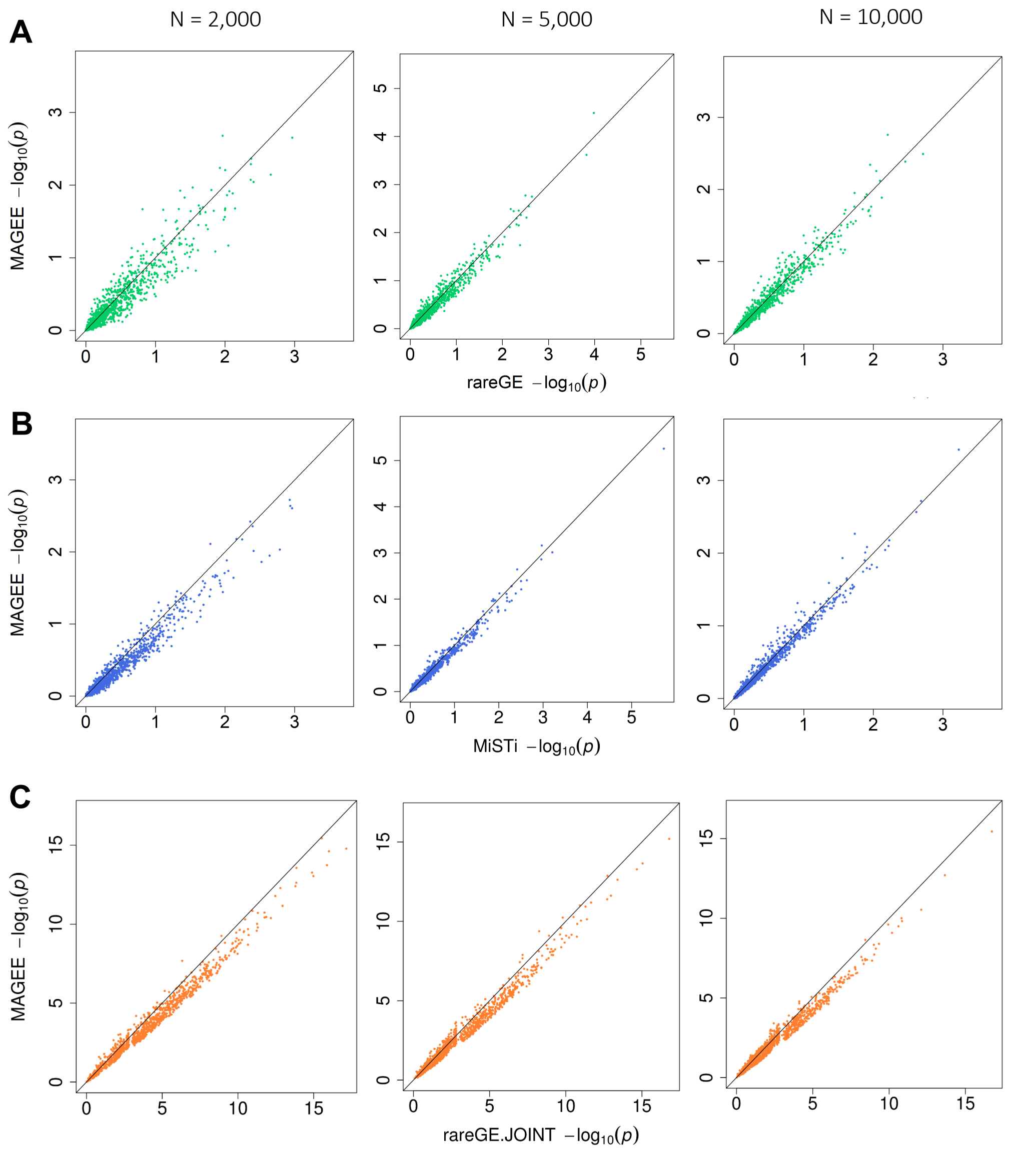

Figure S6. CPU time per $p$ value of MAGEE, rareGE and MiSTi tests on binary traits in unrelated samples. (A) MAGEE, rareGE and MiSTi GEI tests. (B) MAGEE and rareGE joint tests.

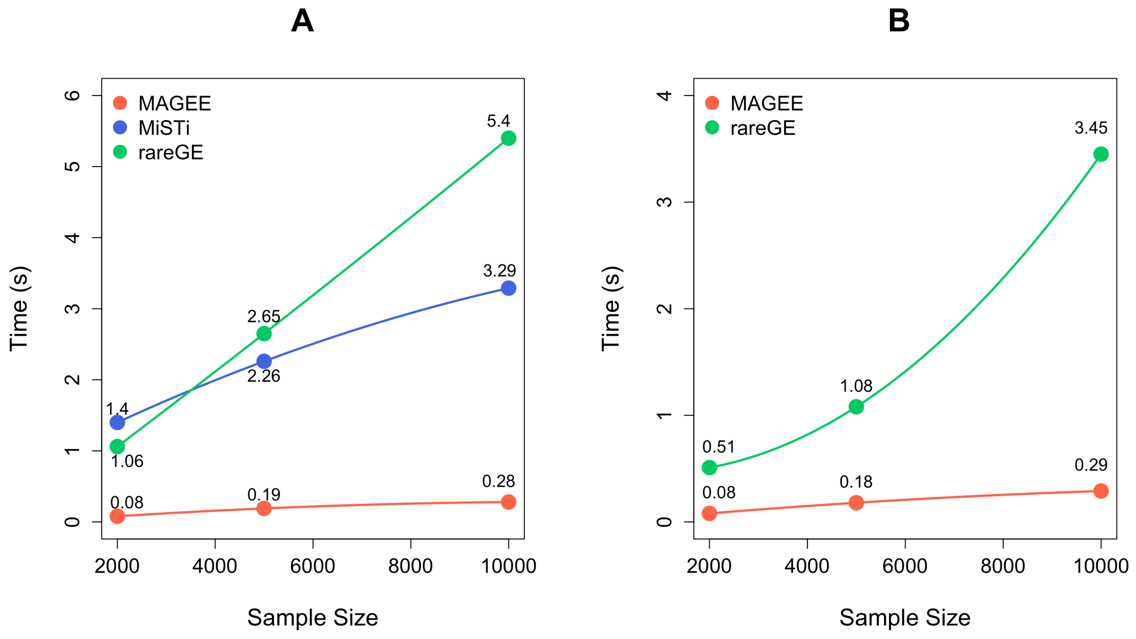

Figure S7. Empirical power of MAGEE tests on binary traits in 20,000, 50,000, and 100,000 related samples. (A) Scenario 1: 80% null variants, 10% causal variants with positive effects and 10% causal variants with negative effects for both genetic main effects and GEI effects. (B) Scenario 2: 80% null variants, 10% causal variants with positive effects and 10% causal variants with negative effects for genetic main effects only. (C) Scenario 3: 80% null variants, 10% causal variants with positive effects and 10% causal variants with negative effects for GEI effects only. (D) Scenario 4: 80% null variants, 16% causal variants with positive effects and 4% causal variants with negative effects for both genetic main effects and GEI effects. (E) Scenario 5: 80% null variants, 16% causal variants with positive effects and 4% causal variants with negative effects for genetic main effects only. (F) Scenario 6: 80% null variants, 16% causal variants with positive effects and 4% causal variants with negative effects for GEI effects only.

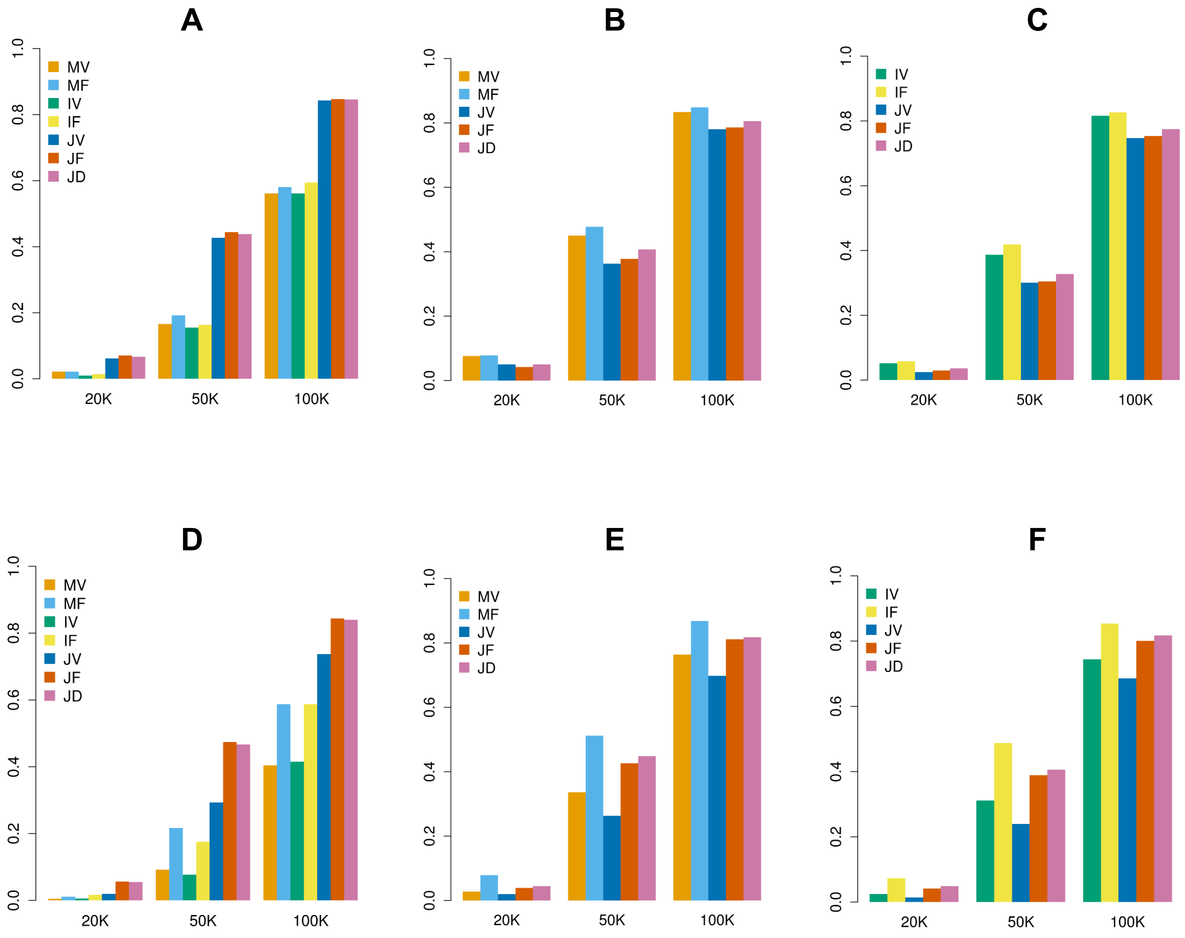
A
B
C
D
E
F
MV
MF
IV
IF
JV
JF
JD
20K
50K
100K
0.0
0.2
0.4
0.6
0.8
1.0

#### TABLES

Table S1. A summary of the test statistics and their distributions under the null hypothesis, and the combination methods of $p$ values for the GEI and joint tests within the MAGEE framework.

| Test statistic | Distribution | Combination Method |
|---|---|---|
| GEI tests | | |
| 1. Interaction variance component (IV) test | | |
| $T_\gamma = \tilde{\boldsymbol{S}}_{\boldsymbol{K}}^T \boldsymbol{W}_{\boldsymbol{K}} \boldsymbol{W}_{\boldsymbol{K}} \tilde{\boldsymbol{S}}_{\boldsymbol{K}}$ | $\sum_{j=1}^{cq} \xi_{\gamma,j}\, \chi_{1,j}^2$, and $\xi_{\gamma,j}$ are the eigenvalues of $\boldsymbol{W}_{\boldsymbol{K}} \boldsymbol{\Lambda} \boldsymbol{W}_{\boldsymbol{K}}$ | None |
| 2. Interaction hybrid test using Fisher's (IF) method | | $p_{IF} = \mathrm{P}(\chi_4^2 > -2logp_{\gamma_0} - 2logp_\tau)$ |
| $T_{\gamma_0} = \tilde{\boldsymbol{S}}_{\boldsymbol{KB}}^2$ | $\xi_{\gamma_0}\chi_1^2$, and $\xi_{\gamma_0} = \boldsymbol{1}_{\boldsymbol{cq}}^T \boldsymbol{W}_{\boldsymbol{K}} \boldsymbol{\Lambda} \boldsymbol{W}_{\boldsymbol{K}} \boldsymbol{1}_{\boldsymbol{cq}}$ | |
| $T_\tau = \tilde{\boldsymbol{S}}_{\boldsymbol{KV}}^T \boldsymbol{W}_{\boldsymbol{K}} \boldsymbol{W}_{\boldsymbol{K}} \tilde{\boldsymbol{S}}_{\boldsymbol{KV}}$ | $\sum_{j=1}^{cq} \xi_{\tau,j}\, \chi_{1,j}^2$, and $\xi_{\tau,j}$ are eigenvalues for $\boldsymbol{W}_{\boldsymbol{K}} \boldsymbol{\Lambda}_{\boldsymbol{KV}} \boldsymbol{W}_{\boldsymbol{K}}$, $\boldsymbol{\Lambda}_{\boldsymbol{KV}} = \boldsymbol{\Lambda} - \boldsymbol{\Lambda}\mathbf{W}_{\mathbf{K}}\boldsymbol{1}_{\boldsymbol{cq}}\left(\boldsymbol{1}_{\boldsymbol{cq}}^T\mathbf{W}_{\mathbf{K}}\boldsymbol{\Lambda}\mathbf{W}_{\mathbf{K}}\boldsymbol{1}_{\boldsymbol{cq}}\right)^{-1}\boldsymbol{1}_{\boldsymbol{cq}}^T\mathbf{W}_{\mathbf{K}}\boldsymbol{\Lambda}$ | |
| Joint tests | | |
| 1. Joint variance component (JV) test | | $p_{JV} = \mathrm{P}(\chi_4^2 > -2logp_{MV} - 2logp_{IV})$ |
| MV test $p$ value $p_{MV}$ | See variance component test SMMAT-S (Chen et al., 2019) | |
| IV test $p$ value $p_{IV}$ | See IV test above | |
| 2. Joint hybrid test using Fisher's (JF) method | | $p_{JF} = \mathrm{P}(\chi_8^2 > -2logp_B - 2logp_{AS} - 2logp_{\gamma_0} - 2logp_\tau)$ |
| MF burden test $p$ value $p_B$ | See burden test in SMMAT-E (Chen et al., 2019) | |
| MF adjusted SKAT test $p$ value $p_{AS}$ | See adjusted SKAT in SMMAT-E (Chen et al., 2019) | |
| $P$ value $p_{\gamma_0}$ for $T_{\gamma_0}$ in IF test | See $T_{\gamma_0}$ in IF test above | |
| $P$ value $p_\tau$ for $T_\tau$ in IF test | See $T_\tau$ in IF test above | |
| 3. Joint hybrid test using double Fisher's (JD) procedures | | |
| $p_B$, $p_{AS}$, $p_{\gamma_0}$ and $p_\tau$ in JF test | See JF test above | Step 1: separately combine the $p$ values for main effects and GEI effects: $p_{MF} = \mathrm{P}(\chi_4^2 > -2logp_B - 2logp_{AS})$; $p_{IF} = \mathrm{P}(\chi_4^2 > -2logp_{\gamma_0} - 2logp_\tau)$; Step 2: combine the main effects MF test $p$ value $p_{MF}$ and GEI effects IF test $p$ value $p_{IF}$: $p_{JD} = \mathrm{P}(\chi_4^2 > -2logp_{MF} - 2logp_{IF})$. |

Table S2. A summary of the simulation scenarios and values of constant $c$ for $p$ value comparison of MAGEE, rareGE, and MiSTi tests in unrelated samples.

| | | Quantitative trait | | Binary trait | |
|---|---|---|---|---|---|
| | | $\beta_t$ | $\gamma_t$ | $\beta_t$ | $\gamma_t$ |
| Scenario 1: +/0/-: 10%/80%/10% for both main and GEI effects | | | | | |
| Sample size | 2,000 | 0.015 | 0.042 | 0.08 | 0.09 |
| | 5,000 | 0.015 | 0.026 | 0.06 | 0.065 |
| | 10,000 | 0.015 | 0.017 | 0.06 | 0.04 |
| Scenario 2: +/0/-: 10%/80%/10% for main effects only | | | | | |
| Sample size | 2,000 | 0.09 | 0 | 0.22 | 0 |
| | 5,000 | 0.06 | 0 | 0.13 | 0 |
| | 10,000 | 0.038 | 0 | 0.082 | 0 |

+/0/-: proportions of variants with positive, null and negative effects.

Table S3. A summary of the simulation scenarios and values of constant $c$ for power comparison of all the MAGEE tests in related samples.

| Quantitative trait | | Binary trait | |
|---|---|---|---|
| $\beta_t$ | $\gamma_t$ | $\beta_t$ | $\gamma_t$ |
| Scenario 1: +/0/-: 10%/80%/10% for both main and GEI effects | | | |
| 0.019 | 0.0094 | 0.019 | 0.0094 |
| Scenario 2: +/0/-: 10%/80%/10% for main effects only | | | |
| 0.025 | 0 | 0.025 | 0 |
| Scenario 3: +/0/-: 10%/80%/10% for GEI effects only | | | |
| 0 | 0.0116 | 0 | 0.0116 |
| Scenario 4: +/0/-: 16%/80%/4% for both main and GEI effects | | | |
| 0.015 | 0.007 | 0.015 | 0.007 |
| Scenario 5: +/0/-: 16%/80%/4% for main effects only | | | |
| 0.02 | 0 | 0.02 | 0 |
| Scenario 6: +/0/-: 16%/80%/4% for GEI effects only | | | |
| 0 | 0.009 | 0 | 0.009 |

+/0/-: proportions of variants with positive, null and negative effects.
